# A supermatrix phylogeny of the world’s bees (Hymenoptera: Anthophila)

**DOI:** 10.1101/2023.06.16.545281

**Authors:** Patricia Henríquez-Piskulich, Andrew F. Hugall, Devi Stuart-Fox

## Abstract

The increasing availability of large phylogenies has provided new opportunities to study the evolution of species traits, their origins and diversification, and biogeography; yet, with the exception of butterflies, taxonomically well-curated phylogenies are currently lacking for major insect groups. Bees (Hymenoptera: Anthophila) are a large group of insect pollinators that have a worldwide distribution, and a wide variation in ecology, morphology, and life-history traits, including sociality. For these reasons, as well as their major economic importance as pollinators, numerous molecular phylogenetic studies of relationships between and/or within families or genera for this group have been published. We used publicly available sequence data, a family-level phylogenomic backbone, and ultra-conserved element (UCE) data, reconciled to a taxonomic database, to produce a dated phylogeny for bees. The phylogeny comprises 4651 bee species, representing 23% of species and 86% of genera. At family, subfamily, and tribe levels, the data were robust, but between and within some genera relationships remain uncertain. In addition, most of the species with available sequence data are geographically distributed in North America and Europe, highlighting gaps that should be considered in future research to improve our understanding of bee evolution and phylogeography. We provide a summary of the current state of molecular data available and its gaps, and discuss the advantages and limitations of this bee supermatrix phylogeny (available online at beetreeoflife.org), which may enable new insights into long standing questions about evolutionary drivers in bees, and potentially insects.

**Highlights:**

- Bee supermatrix phylogeny constructed with public and published sequence data.
- Includes 23% of currently recognised species and covers 86% of genera.
- Provides a summary of remaining gaps in bee phylogenetics.
- Available online at beetreeoflife.org, with subsetting tool to facilitate comparative analyses.

## 1. Introduction

Phylogenies are a key tool, not only for evolutionary biology, but also ecology subdisciplines such as conservation biology, community ecology and macroecology (Beck et al., 2012; Harvey and Pagel, 1991; Mouquet et al., 2012). In the last two decades, phylogenetic supermatrices of several groups have become available, opening new opportunities to study the evolution of species traits, their origins and diversification, and biogeography. Within the animal kingdom, these supermatrix phylogenies comprise mainly vertebrate groups such as marsupials, bats, passerine birds, gobies, and cetaceans, to name a few (Amador et al., 2018; Jønsson et al., 2016; McCraney et al., 2020; McGowen et al., 2009; Mitchell et al., 2014). Despite insects constituting most of the species on earth (Mora et al., 2011), well-curated large scale phylogenies are currently lacking for major insect groups, with the exception of butterflies (Adler and Foottit, 2018; Chazot et al., 2021; Kawahara et al., 2023).

Bees (Hymenoptera: Anthophila) are a large and diverse group of insect pollinators with over 20000 described species and a worldwide distribution (Orr et al., 2020). This group has received increased attention in recent decades given their importance as pollinators and the future consequences of insect decline (Potts et al., 2016; Wagner, 2020; Winfree et al., 2009). Bees also hold special interest for understanding life-history evolution (Michener, 2007). Species broadly fall into one of three life-history categories (Danforth et al., 2019): i. Solitary, comprising more than 75% of described species where all females are able to produce offspring and each build and provision their own nest; ii. Social, representing little less than 10%; where species display division of labour, cooperative brood care and generational overlap; iii. Parasitic, including close to 13% of species, where most are brood parasites and do not build a nest or forage for food and instead lay their eggs in the nest of other bee species, and a few are social parasites where the female replaces the queen, takes over the colony and co-opts worker females to rear her offspring. For some species the lines between these categories are blurred, mainly in the bee family Halictidae, where some species are socially polymorphic (Davison and Field, 2018; McFrederick et al., 2014; Plateaux-Quénu et al., 2000; Richards et al., 2003). Bee species also differ in their morphological features, annual life cycle, diet breadth, and nesting behaviour. The latter includes a variety of substrates such as soil, wood, or pithy stems, and the use of diverse materials including their own glandular secretions, and a wide range of foreign resources (e.g., floral oils, leaves, resin, sand, pebbles) (Michener, 2007). The diverse life history and natural history of this remarkable group of insects make them ideal for macroecological and macroevolutionary analyses.

The first attempt to produce a supermatrix phylogeny of bees was by Hedtke et al. (2013) a decade ago. Since then, the amount of data that has become available has increased dramatically, demanding a substantially updated phylogeny for this group. Numerous molecular phylogenetic studies of relationships between and/or within families or genera for this group have been published over the years (Supplementary Table S1). In addition, comprehensive molecular phylogenies have been published for the seven bee families (Melittidae, Andrenidae, Halictidae, Colletidae, Stenotritidae, Megachilidae and Apidae (Michener, 2007)). Although some uncertainties remain unresolved, the estimates of most of their evolutionary relationships between families and subfamilies are robust (Almeida et al., 2019, 2012; Bossert et al., 2021, 2019; Danforth et al., 2008; Gonzalez et al., 2012; Litman et al., 2011; Michez et al., 2009; Peters et al., 2011; Sann et al., 2018). As a result of this volume of phylogenetic studies, a substantial amount of phylogenetic data has become available for bees which can be used to build a supermatrix phylogeny.

Here, we use publicly available sequence data, a family-level transcriptome phylogenomic backbone, and ultra-conserved elements (UCE), reconciled to a taxonomic database to ensure taxonomic consistency, to construct a supermatrix phylogeny of bees. We focused on curating and analysing widely sampled loci used in previously published phylogenies to maximize overlap of data among lineages and reduce supermatrix sparseness. Once all components were curated, we used them to produce a concatenated supermatrix to construct maximum likelihood trees. This approach produced the largest bee phylogeny to date that includes 4651 species in 427 genera. While this supermatrix phylogeny doesn’t aim to solve current uncertainties in phylogenetic relationships, it is a curated composite of extensive previous phylogenetic work which may enable multiple new research opportunities. In addition, it provides information regarding data gaps that need to be addressed to improve resolution for some bee genera. This supermatrix is available for download and can be subsetted through the online tool beetreeoflife.org. In addition, the website also hosts 1000 bootstrap sample trees converted to dated chronograms to represent the phylogenetic information and uncertainty in our supermatrix.

## 2. Methods

### 2.1. Taxonomic database

Large scale phylogenies require a consistent taxonomy and nomenclature to which all data can be reconciled (e.g., Driskell et al., 2004; Hosner et al., 2022; Thomas et al., 2013). For this work, a taxonomic database was developed for Anthophila based on data provided by Orr et al. (2020), which included synonyms, and a more accurate and curated version of current nomenclature than the Open Tree of Life project (Rees and Cranston, 2017). We modified this database to reflect revised generic-level classification from more recent published taxonomic references that have provided generic stability. This taxonomic database only includes species that are part of the phylogeny, recognises seven bee families and 28 subfamilies, and provides a list of genera, species, and notes on nomenclature decisions and the taxonomic research that supports them (Supplementary file 1). In addition, differences exist regarding nomenclature between some databases (e.g., Integrated Taxonomic Information System, Catálogo Moure para as espécies de abelhas neotropicais, Atlas Hymenoptera, Catalogue of Life), and thus the taxonomic database also includes alternate names for those species with names that have not been updated to follow the most recent nomenclature decisions. Because taxonomy and nomenclature are constantly changing, especially for taxa where relationships are yet to be resolved, the goal of this taxonomic database is to make it easier for researchers to track nomenclature changes in the phylogeny and easily change them if necessary.

### 2.2. Supermatrix data components

We selected the widely-used supermatrix approach (Chesters, 2020; Driskell et al., 2004; Hedtke et al., 2013; Jetz et al., 2012; Upham et al., 2019) to build a large bee ‘tree-of-life’ over other methods (e.g., supertree: Bininda-Emonds, 2004; Sanderson et al., 1998). Supermatrix methods use all available information to estimate relationships which provides higher resolution, simultaneously providing branch length and uncertainty (bootstrap resampling). We built our phylogenetic supermatrix database for bees based on four key components: i. gene fragment accessions downloaded from global sequence databases; ii. a family-level backbone using a subset of transcriptome phylogenomic data; iii. published UCE datasets combined and condensed; and iv. nomenclature reconciled to the taxonomic database.

Generally, efficient supermatrices have a hierarchical mix of ‘fast’ and ‘slow’ genes, which typically involves a mitochondrial gene for the tips and a set of nuclear loci that can be limited to representatives of the major lineages (Beaulieu and O’Meara, 2018; Driskell et al., 2004). Thus, instead of sheer data volume, we focussed on fewer widely sampled loci used in published studies with the greatest proportion of taxa and phylogenetic spread to maximize overlap of data among lineages. This approach can make it easier to check individual genes, and limit the proportion of taxa without data in common, which can distort phylogenetic inference (Freyman, 2015; Sanderson et al., 2010; Wiens and Morrill, 2011). Thus, the publicly available gene fragment accessions provided the species (tips), the transcriptome phylogenomic data provided a robust backbone consistent with phylogenomic information on tree shape (topology, relative branch length and outgroup rooting), and the UCE leveraged new and powerful phylogenomic data that is increasingly being used, to further enforce best-estimate tree shape. Collation and curation of each component was done with custom Bash scripts and Microsoft Excel (see Supplementary file 2 and Supplementary file 3) (Microsoft Corporation, 2021); details for each component are given in the following sections. Given the multiple data sources, bespoke taxonomic database, and the need for hands-on refinement, we expanded our supermatrix assembly system used in previous works (e.g., Hugall and Stuart-Fox, 2012; Oliver et al., 2023).

#### 2.2.1. Gene fragment accessions

Data were downloaded from the NCBI nucleotide collection (https://www.ncbi.nlm.nih.gov/genbank). We used gene sequences obtained from key phylogenetic references in BLAST+organism searches with an E-value threshold of 5e-6 and retained aligned regions, excluding model and environmental data. For protein coding genes and ribosomal RNA genes we used tblastn and blastn respectively. In total, we obtained data for seven nuclear protein-coding genes (ArgK, CAD, NaK, Pol II, Wnt-1, LW Rh, and EF-1α), two ribosomal genes (16S and 28S), and two mitochondrial coding genes (Cytb and COI) (Table 1).

**Table 1.**
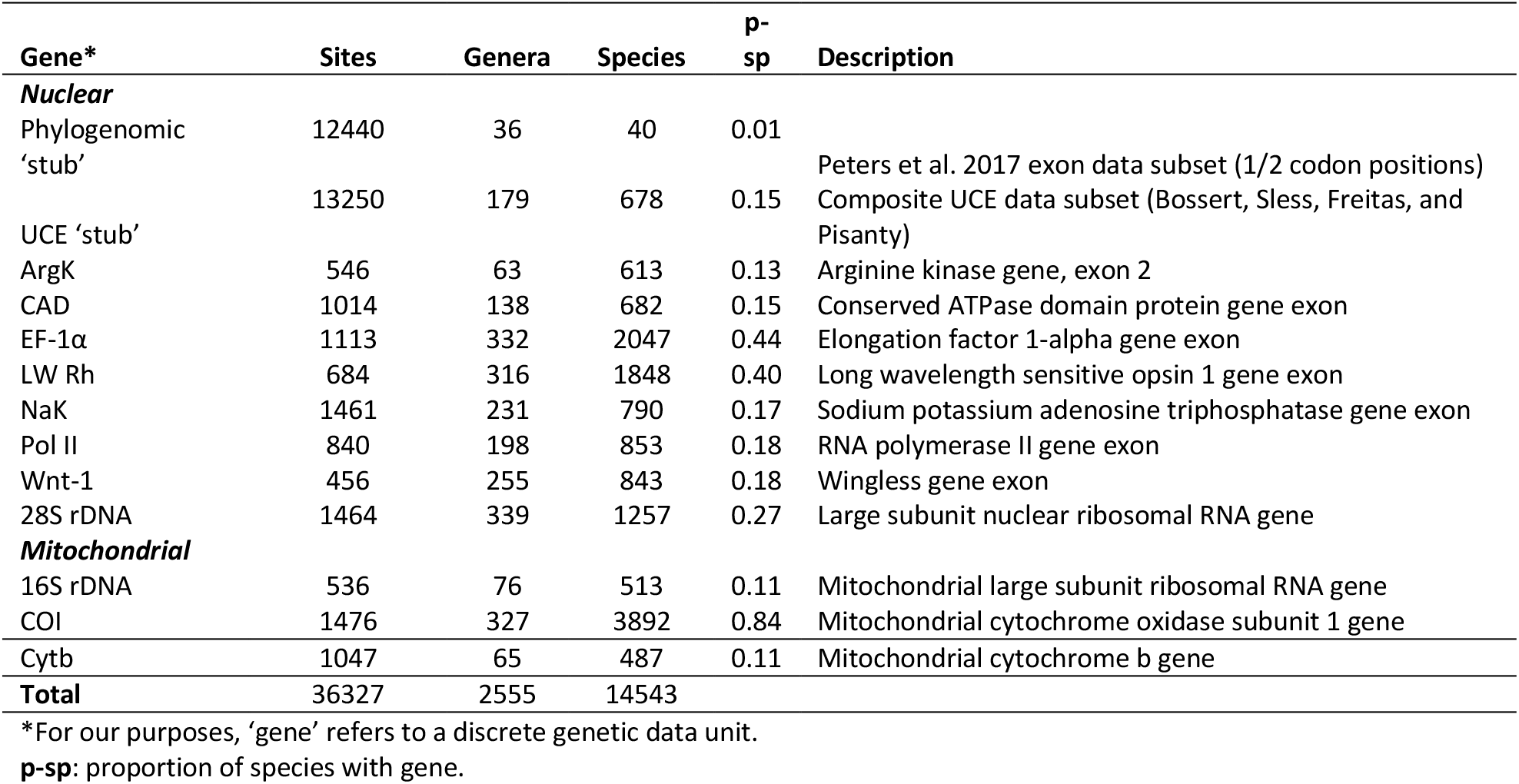
Sampling summary across genetic data.

We focused on protein coding regions (e.g., exons), as they are easier to align, and make it simple to add new data. For the purposes of this work, we did not include the widely sequenced ribosomal 18S gene due to its low and poor phylogenetic signal (Soltis et al., 1999). Because of the limited coverage of whole mitogenome data for bees (Husemann et al., 2021), we extracted only near full-length mitochondrial genes (16S, Cytb, COI) with the reference BLAST. Rather than selecting one ‘best’ longest species accession per locus, we retained all accessions to first assess the gene tree monophyly of nominate species as well as genera. Due to the great number of accessions, EF-1α, LW Rh and COI genes were split into two sets: Apidae and the rest.

Gene accessions were aligned with MAFFT v. 7.245 (Katoh and Standley, 2013) using the auto setting. We then assembled alignments into a custom Microsoft Excel database, and reconciled nomenclature with the taxonomic list (Supplementary file 1). We refined protein coding gene alignments by hand in BioEdit (Hall, 1999) using the original references as a guide, to have all coding regions strictly in-frame. Despite being time consuming, aligning protein coding genes this way makes adding taxa simpler. For the ribosomal RNA genes, we used the MAFFT alignment as is.

We inferred gene trees through IQ-TREE (Nguyen et al., 2015), and in some cases RAxML (Stamatakis, 2014). IQ-TREE used ultrafast bootstrap consensus (Hoang et al., 2018) with models of sequence evolution identified by ModelFinder implemented in IQ-TREE (Kalyaanamoorthy et al., 2017). RAxML used fast bootstrap and the GTR+G model, with the standard -f a setting implemented on CIPRES (Miller et al., 2010). We assessed gene trees for non-monophyletic genera and species, and aberrant sequences such as paralogs (e.g., EF-1α and LW Rh variants) and gross miss-alignments (long terminal branches). Then, we updated the dataset by removing abnormal accessions and in some cases revising the nomenclature. Datasets were realigned if necessary.

The mitochondrial COI was processed slightly differently because of the extensive number of accessions that are publicly available, its nominal intra-specific diversity, and because it is often sequenced in separate fragments. We reduced this gene alignment to a single consensus sequence per species, based on the most common base per site (with ties scored as ambiguous). This approach tends towards the most commonly sequenced sub-lineage and is a simple way to combine data, discount rarer aberrant sequences and rationalize choice of intra-specific lineage complexity. The consensus alignment was then subjected to the same procedure of gene tree and genera monophyly assessment as described for the other genes included in this work. Additionally, we included COI data from The Barcode of Life Data System (BOLD, http://www.boldsystems.org) for a selection of species that had multiple samples that clustered closely (>5%), or that represented taxa we already had with other genes if there was only one sequence available, and that fell within the lowest rank (genus or subfamily depending on taxon sampling) in our working COI tree. Information on data sources for all taxa and genes is summarized in Supplementary file 3.

#### 2.2.2. Family-level phylogenomic backbone

Hymenoptera transcriptome phylogenomic work of Peters et al. (2017) provided the current best estimate of the higher family level phylogenetic tree for Anthophila. This work included data for 41 bee taxa from six of the seven families (excepting Stenotritidae) for 3256 genes. This vast amount of data was unnecessary for the purposes of the bee supermatrix and thus, we extracted a subset sufficient to reconstruct tree shape. To this aim, we trimmed down the first and second position exon datamatrix from [dataset] Peters et al. (2017) to the Anthophila and key closest ‘Crabronidae’ outgroups (60 taxa), and condensed codons (3.01 million bases, 1.5 million codons) down to a practical amount (12.4 kb). This was done by deleting codons with <85% of taxa and then randomly sub-sampling one third of codons (jack-knife), thereby reducing the proportion of missing data from the original 57.3% complete to 92.8% complete. We then used this phylogenomic ‘stub’ to produce a tree, as previously described for the gene fragment datasets, and compared it with the original Hymenoptera phylogeny by Peters et al. (2017).

#### 2.2.3. UCE data

Ultra-conserved elements are a method of extracting phylogenomic scale data that is increasingly being used for bees. Because this method produces more data than is necessary for the bee supermatrix phylogeny, we selected and combined a subset of data from five previously published data-matrices ([dataset] Bossert, 2021a, 2021b; [dataset] Freitas et al., 2020; [dataset] Pisanty et al., 2022; [dataset] Sless, 2021). First, we produced consensus sequences for each of the five data-matrices with the same approach described for the COI gene. Then the 1388 UCE consensus loci obtained from [dataset] Bossert (2021a) were mapped onto the consensus sequences of the other data-matrices with BLAST v2.13.0, to identify the corresponding orthologous sections. We then selected a subset of suitable UCE loci using several criteria: maximum e-value of 1e-50, >150 sites, and >50% sites matched to the consensus reference. We aligned these loci with MAFFT as previously described for gene fragments, concatenated them into a UCE ‘supermatrix’, and reconciled species names with the taxonomic database. This supermatrix (57.9 kb in 94 loci for 777 samples) was then compacted down to an amount adequate to reconstruct key tree shape, without excess computational burden, by deleting sites that were <74% complete and then further reduced by one third jack-knifing, resulting in a dataset of 13.3 kb; 85% complete. We then produced a tree to compare to the original published UCE phylogenies, and to assess for non-monophyletic genera and aberrant sequences as previously described. The resulting tree had 71% of nodes with bootstrap support >=90%. Details of the primary transcriptome and UCE data sources are provided in Supplementary file 3.

There are three approaches when working with UCE sequence data: i. Reconstruct from primary reads; ii. reconstruct from locus datasets; iii. reconstruct from the final processed and refined supermatrix. We took the third approach, as a simple compromise that has the benefit of leveraging from previous bioinformatic work. We regard it preferable to ‘supertree’ approaches (e.g., Kimball et al., 2019) that merely use the topology of phylogenomic datasets, with branch length/ages applied post-hoc.

### 2.3. Supermatrix assembly and analysis

Once all supermatrix data components were reconciled to the taxonomic database, aligned and assessed for aberrant elements, they were used to produce a concatenated supermatrix. We did this by taking the best (longest accepted) single exemplar sequence per ‘gene’ per species, for all species with a total of least 400 sites of any data. The supermatrix was then compacted by removing regions with little or no data (<10% taxa with any data per ‘gene’) and ambiguous alignment regions with Aliscore v2.2 (Misof and Misof, 2009), while maintaining protein coding reading frames.

Sparse, patchy supermatrices often include taxa groups that share no data in common, which can bias the phylogenetic inference (Sanderson et al., 2010; Smirnov and Warnow, 2021; Wiens and Morrill, 2011). To ameliorate this issue, we implemented minimal taxonomic rank substitution to improve data density on the phylogenomic ‘stub’, and species with no overlap of data within genera (Pennell et al., 2016). For genera that contained species that did not share any data with any other member (i.e., the only species in that rank with that genetic data), we reassigned the data to the member of the genus with the most data and removed the source species. In total 40 substitutions were made (0.4% of the matrix).

The outgroup included five taxa representing the Crabronidae subfamilies Astatinae, Bembicinae, Crabroninae and Philanthinae, based on the phylogenomic ‘stub’ plus a small amount of gene fragment data. As gene data was patchy across outgroups, taxonomic rank-substitution was also used to create composite family representatives for some taxa. The gene composites included the genera *Astata* (Astatinae), *Bembix* (Bembicinae), *Cerceris* (Philanthinae), *Philanthus* (Philanthinae) and *Tachysphex* (Crabroninae).

To reduce computational burden, the supermatrix was split into three subsets comprising the families Megachilidae + Andrenidae + Melittidae, Halictidae + Colletidae + Stenotritidae, and Apidae, each sharing a core set of 18 family representatives with maximal data, and the five outgroups. In addition, we extracted one species with the most data for each nominal genus to create a genus-level supermatrix. For each of these supermatrix subsets, phylogenetic inference was done by IQ-TREE with ultrafast bootstrap, using a RAxML starting tree, all nearest-neighbour interchange and refined tree search settings (-allnni, -beps 8, -bnni, -bcor 0.98, -wbtl, -pers 0.4, -nstop 200, --sprrad 8; via CIPRES). The intention was to focus effort on tree support space, in what can be a difficult problem (Shen et al., 2020). The optimal partitioned sequence evolution model was determined with ModelFinder (in IQ-Tree) using the genus-level supermatrix, and this 8-partition model scheme was then used for all remaining analyses (see Supplementary file 3). This ensures a common model irrespective of the amount of data per gene per family subset (i.e., as if all three were analysed simultaneously). The trees from the three family subsets were then joined together again using the common family representatives, for both the consensus tree and the 1000 bootstrap trees.

### 2.4. Tree dating calibration

Patchy and missing data can exacerbate unequal/uneven estimates of branch length among lineages (how clock-like the tree appears), over and above inherent rate variation (Roure et al., 2013; Zheng and Wiens, 2015). To reduce this calibration problem, it is necessary to rely on common partition models, and relaxed-clock models. The species-level supermatrices were far too large to properly run Bayesian relaxed-clock methods (Fisher et al., 2022), and the divide and graft approach is complicated and constraining (e.g., Jetz et al., 2012; Upham et al., 2019). Thus, to overcome these issues we sought a simpler approach, explained in the following paragraphs, that balanced computational effort with the limited precision of the data.

Trees were converted into chronograms using Penalised Likelihood Rate Smoothing (PLRS) (Sanderson, 2002), using treePL (Smith and O’Meara, 2012). To enforce chronograms to remain fully bifurcating and improve the efficiency of the PLRS iteration, a small length increment was added to any branch of length >0.0005 (amounting to mean 0.43% of the original total tree length, affecting 6.8% of branches). TreePL was run with ‘thorough prime’ optimized settings and smoothing factor=10. Because the all-species tree was too large to use cross-validation, a suitable smoothing factor was empirically determined from the genus-level tree.

There are few certain dating calibrations for major lineages within bees and considerable variation among published works (e.g., Almeida et al., 2012; Cardinal and Danforth, 2013; Freitas et al., 2022; Peters et al., 2017; Schwarz et al., 2006). Many studies rely on secondary dating from a limited number of previous works (e.g., Martins et al., 2014; Rehan and Schwarz, 2015; Trunz et al., 2016). Therefore, rather than enforce a range of putative calibrations, we simply calibrated the bee root to a broad normal distribution of 100 million years ago (mya) SD 4. This was achieved by drawing a root age from a normal distribution for each of the 1000 bootstrap replicates. This was used for both the genus-level and the all-species trees. The results were then compared to published ages/calibrations for a suite of key higher clades.

## 3. Results and discussion

Our intention was to produce a useful resource, by collating molecular data for bees, summarizing the current state of the data, and providing a serviceable large-scale supermatrix phylogeny. Molecular data is continuously being generated, and in this process higher taxonomy is also being reconciled. As a result, most of the higher ranks are consistent and well supported: family, subfamily, tribe; but to a lesser extent, genera. We kept the tree inference to an efficient approximation, with appropriate computational effort to match the level of precision inherent in the data. This phylogeny is not a test of bee systematics but a reflection of current information and ideas about bee relationships. In the following sections we provide information regarding the current state of the data, the estimated phylogeny, and the uncertainty for the set of phylogenetic trees produced in this work.

The phylogeny includes 4651 species and 427 genera, which represent 23% of currently described species and 82% of genera, respectively (Fig. 1). In addition, the genera included contain 96% of all described species. This comprises 36327 sites with a total of 18.6 million base characters from 14543 data elements, median 11% complete (3981 sites per species). Supplementary file 3 provides complete information on genetic data for the entire supermatrix. We took a more hands-on approach to curating suitable data rather than an all-in automation to build our dataset (Beaulieu and O’Meara, 2018), with several iterations of collation and summary inference guiding its development. Our dataset is not exhaustive but a substantial representation of the total possible molecular data, as of April 2023, providing the largest supermatrix dataset and tree of bees to date (Chesters, 2020; Hedtke et al., 2013). Tables 1, 2 and Supplementary Table S2 show brief summaries of our sampling by family and by gene. A more detailed breakdown by subfamily is provided in Supplementary Table S3. Between families, sampling coverage of named genera ranged from 73-100% but at the species level, coverage was lower ranging between 19-41% (Table 2). Taxonomic sampling is key for phylogenetic inference accuracy (Nabhan and Sarkar, 2012), and while family, subfamily and tribe level relationships are robust, further work is necessary to fill in the gaps and reduce uncertainties in relationships between and within genera.

**Fig. 1.**
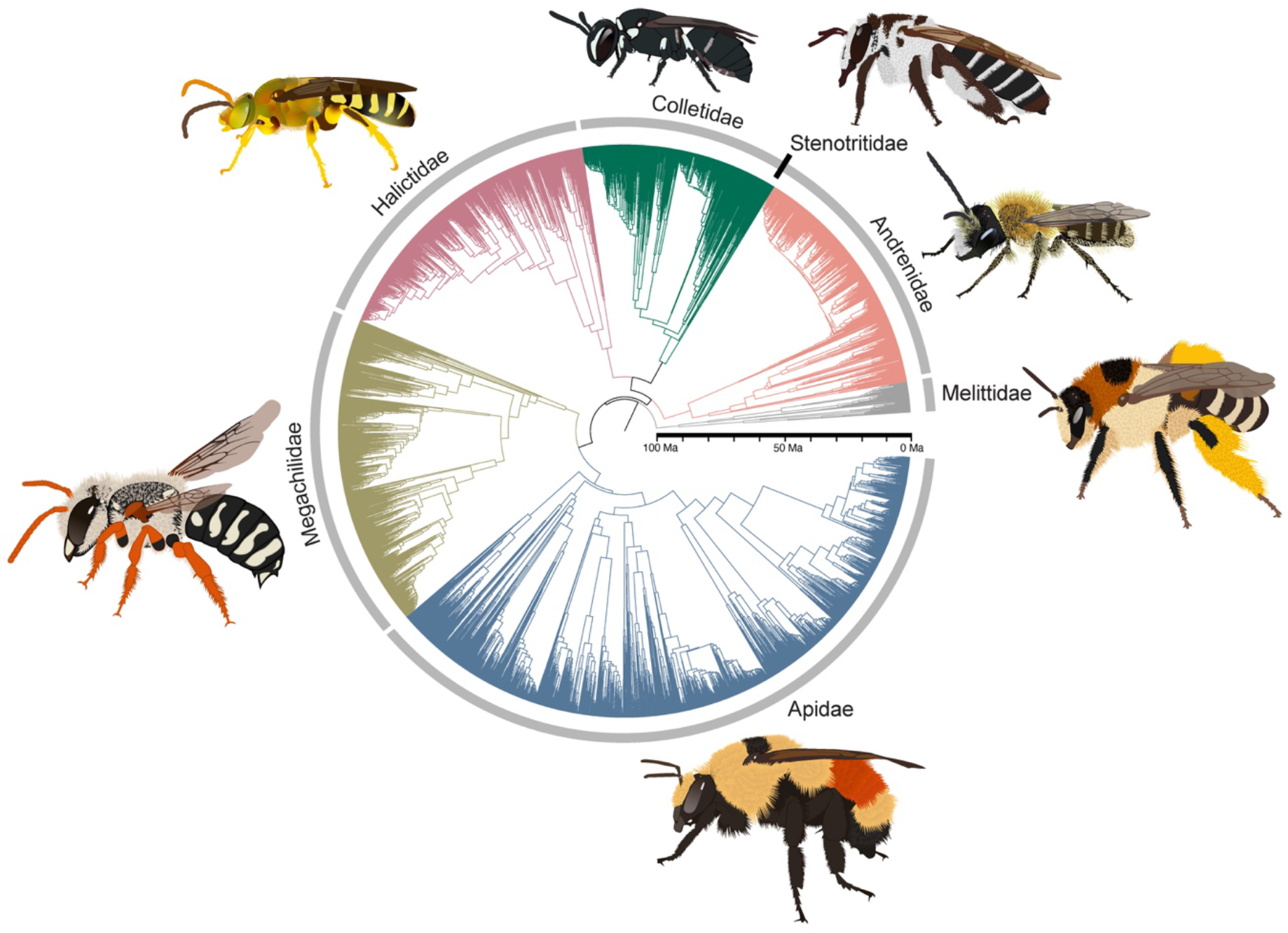
Supermatrix phylogeny for Anthophila built with public and published sequence data. The full tree is available for download at beetreeoflife.org and includes 4651 species in 427 genera, including the species illustrated in the figure: *Agapostemon texanus* (Halictidae), *Andrena fulva* (Andrenidae), *Anthidium chilense* (Megachilidae), *Bombus huntii* (Apidae), *Ctenocolletes nigricans* (Stenotritidae), *Dasypoda hirtipes* (Melittidae), and *Hylaeus affinis* (Colletidae). Family colour coding used throughout.

**Table 2.** Summary of taxonomic sampling by family.

| Family | Total subfamilies* | Total genera | Included genera (%) | Total species | Included species (%) |
| --- | --- | --- | --- | --- | --- |
| Andrenidae | 3 | 50 | 43 (86) | 2957 | 644 (22) |
| Apidae | 5 | 211 | 184 (87) | 5829 | 1779 (31) |
| Colletidae | 9 | 76 | 64 (84) | 2616 | 523 (20) |
| Halictidae | 4 | 82 | 62 (76) | 4400 | 779 (18) |
| Megachilidae | 3 | 83 | 61 (73) | 4099 | 838 (20) |
| Melittidae | 3 | 15 | 11 (73) | 201 | 82 (41) |
| Stenotritidae | 1 | 2 | 2 (100) | 21 | 6 (29) |
| **Total** | 28 | 519 | 427 (82) | 20123 | 4651 (23) |

\*All subfamilies are included in this work and total numbers for each rank are based on our revised taxonomic list.

### 3.1. Geographic distribution of sampling

Species in our tree are biased towards the Neartic, followed by the Paleartic and Neotropics (Fig. 2, Supplementary Table S4), indicating knowledge gaps in bee distribution and under-sampling of all other regions (see Supplementary file 4 for a description of the methods used to obtain bee distribution data). Despite bees being a diverse insect group with global distribution (Danforth et al., 2019; Michener, 2007), geographical biases remain. This has previously been discussed in terms of ecological gaps (Archer et al., 2014), and taxonomic gaps (Orr et al., 2020), but efforts should also be made to obtain molecular data from other regions to better understand the relationships and the evolution of bees, especially considering that some of these less studied regions might host high bee species richness (Batley and Hogendoorn, 2009; Eardley et al., 2009; Freitas et al., 2009; Groom and Schwarz, 2011; Melin and Colville, 2019).

**Fig. 2.**
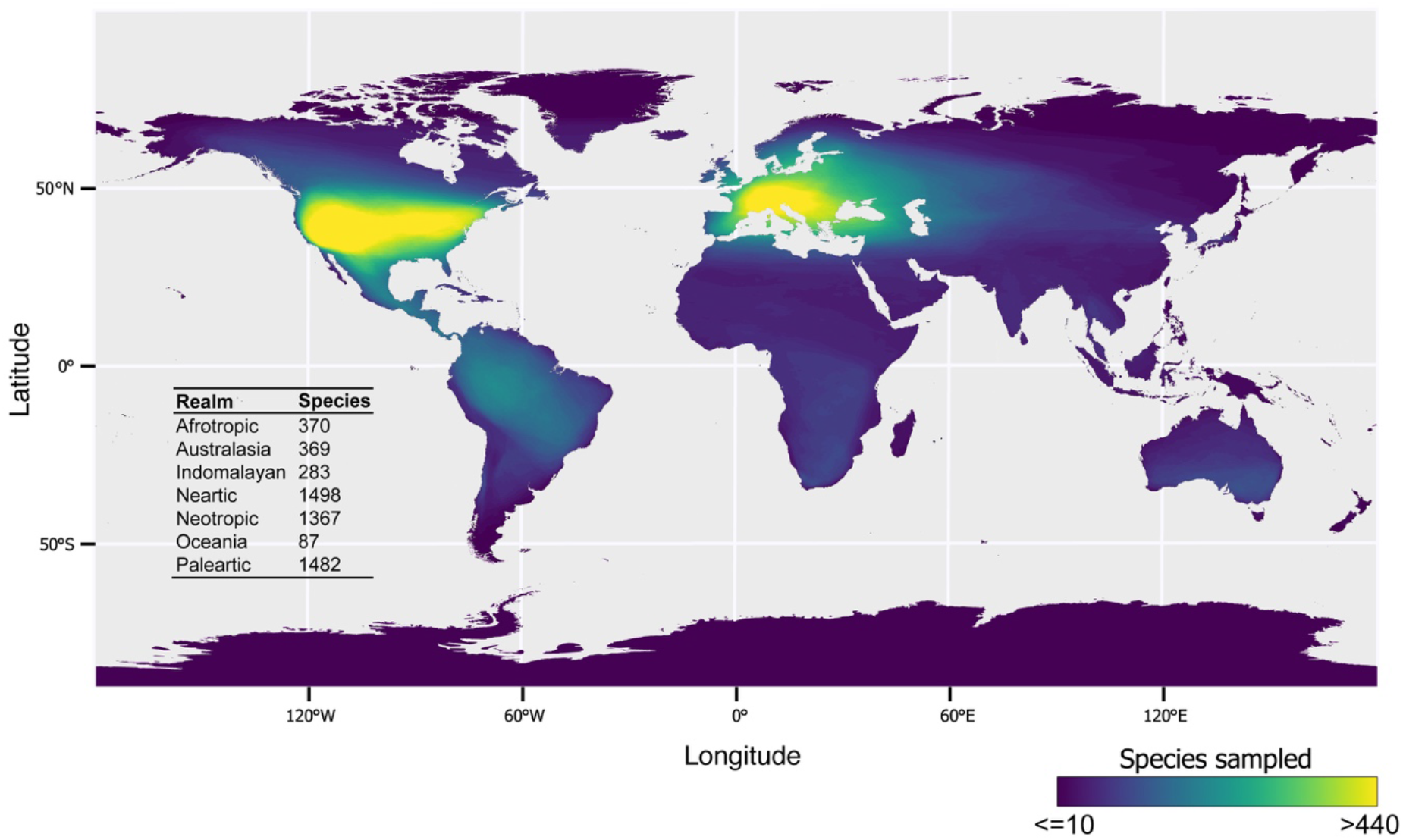
Distribution of bees with available molecular data that was used to build the supermatrix phylogeny.

### 3.2. Gene sampling

Overall, the data were highly skewed with many species (38%) having one gene and only a small number (16%) having more than five (Fig. 3A-B). However, data were well distributed across linages and a small number of species with many genes were phylogenetically spread across the tree (Fig. 3A-C). In addition, the use of shortened subsets of the very large phylogenomic data (transcriptome and UCE) provided sufficient information to ensure the tree was consistent with published higher-level phylogenies in topology and relative branch length. Our phylogenomic ‘stub’ recreated published phylogenomic trees with high bootstrap support (>95%), and the UCE ensured that most of the key nodes matched previously published results (see Similarity to previous studies section). Thus, the addition of thousands of taxa with little data to the core backbone had limited effect on the underlying backbone of the tree. This can be seen in the similarity of the genus representative tree and the equivalent subtree pruned from the full all-species tree where 89% of nodes were recovered; remaining nodes had low support in either or both trees (Fig. S1). Branch lengths were also very similar as are the inferred clade ages (see section below).

**Fig. 3.**
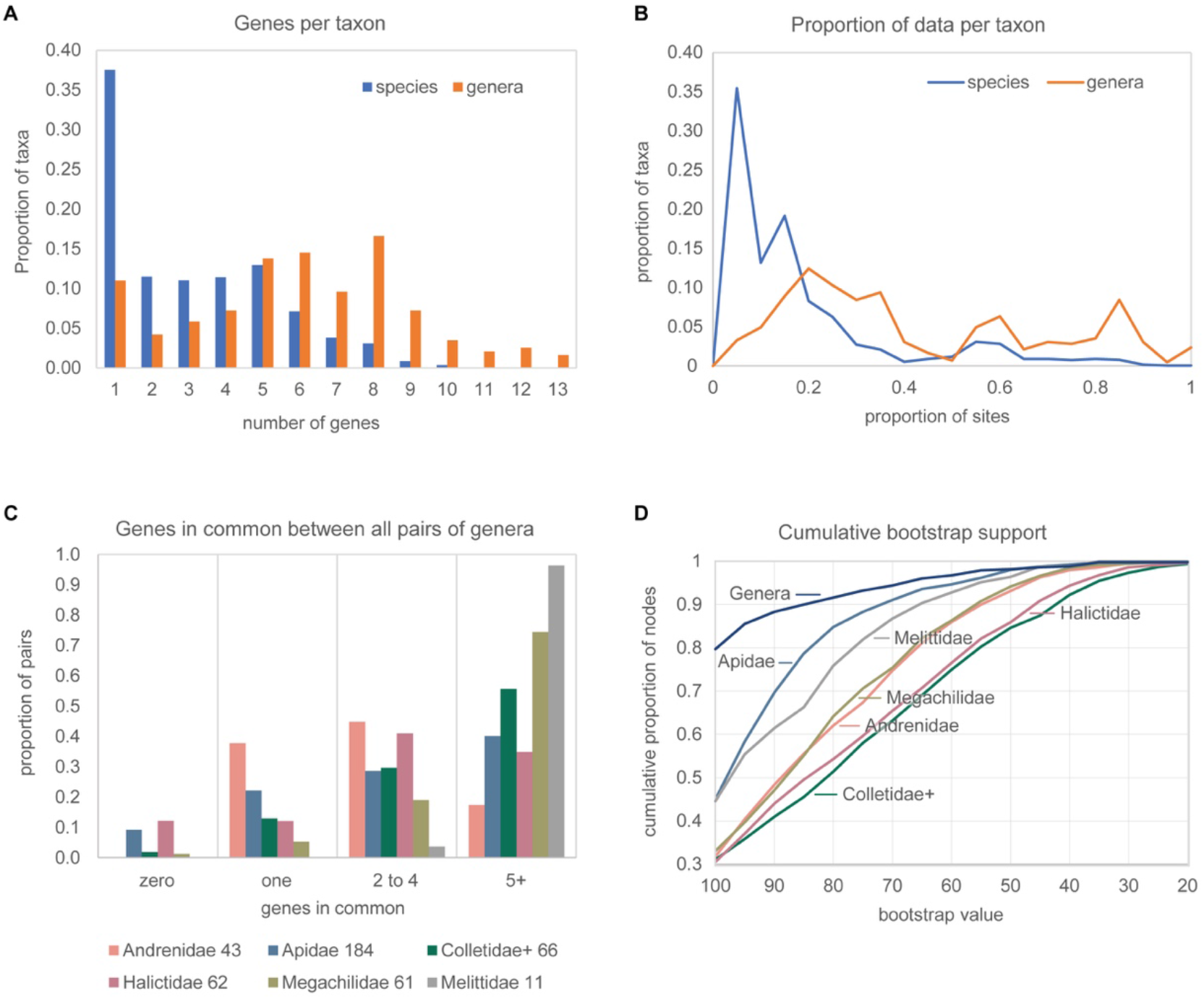
Graphical summaries of data completeness and tree support. **A.** Number of genetic data elements (‘genes’) per species in the 4651 species tree and per genus in the 427 genera representative tree. **B.** Proportion of sites with data in the supermatrix not including the phylogenomic stub. **C.** Number of genes common between all genera within each family, summarized in four categories (number of genera after name, and in histogram left to right in alphabetical order). **D.** Cumulative bootstrap support across nodes by family and the genus representative tree. Colletidae+ includes Stenotritidae.

Based on cumulative IQTree ultrafast bootstrap support values (ufbs) (Fig. 3D), median ufbs support was 89, while 23% of nodes were below 70. This varied across the family subtrees. The Apidae and Melittidae had the best support (median ufbs 94 for both), and Colletidae the least (median 77). The genus-level representative tree returned higher support: median ufbs 95 (Fig. 3D, Fig.S1).

### 3.3. Data structure limitations

A key issue in supermatrix data structure is how much data is shared between true sister taxa, a simple measure being the genes taxa have in common (hereafter referred to as ‘GIC’ – genes in common). GIC is very similar to taxon triplets measures (Sanderson et al., 2010), and provide an indication of data structure limitations by identifying groups that are most affected by scarce data overlap, and would therefore profit from targeted data gap filling (see Freyman, 2015). In a cladistic analysis, taxa with no data in common cannot be recovered as sister taxa (Sanderson et al., 2010). In principle, a supermatrix would have at least one GIC to all species (typically mtDNA) with the remaining genes (typically nuclear) limited to key representatives of the major lineages. However, in practice supermatrices tend to be patchy with much missing data, potentially leaving many true sister taxa with little or no data in common. Nonetheless, our most common gene COI covers 3892 species (84%, Table 1). The nuclear genes EF-1a and LW Rh are also widely sampled (44 and 40%, respectively).

At family and subfamily levels, data was highly robust and structured with a minimum of seven GIC (median of eight) among all subfamilies (Supplementary Table S5, Fig. 3C). Within genera the overlap of data varied (Supplementary Table S5, Fig. S2A-B), with numerous cases of zero shared data between individuals within a genus (see Supplementary Table S6 for details). The generally increasing GIC deeper into the tree and variation in GIC towards the tips is apparent from GIC mapped onto the tree (Figure S2C; note that in the tree GIC should be >0). There are two approaches when dealing with limited GIC, either remove species or fill in the existing gaps. Excluding rogue taxa may also help but this can be a complicated process (Smith, 2022). Until these gaps are filled in, species relationships within poorly structured genera should be treated with caution, and in the cases where analyses are sensitive to tree topology, these genera could be pruned back to single genus representatives.

### 3.4. Similarity to previous studies

As intended, our tree largely reproduces the recent best phylogenomic scale results for families and subfamilies (e.g., Bossert et al., 2021, 2019; Branstetter et al., 2017; Peters et al., 2017; Sann et al., 2018; Sless et al., 2022). These differ somewhat to older multi-gene studies, the data of which we also included (Almeida et al., 2012; Cardinal et al., 2018; Cardinal and Danforth, 2013; Ramos et al., 2022; Rehan et al., 2010). There are some differences between our full species tree and our genus representative tree, and these differences mirror the uncertainties reported in some of the primary studies. Most of these differences involve short internodes between lineages. Even with massive amounts of multi-locus UCE data such situations can be unresolvable and/or uninformative (Degnan and Rosenberg, 2009; Hahn and Nakhleh, 2016). Some key remaining uncertainties include the subgeneric relationships within *Andrena* (Andrenidae: Andreninae) given its high degree of paraphyly and polyphyly, where classification changes will likely occur with further efforts to resolve such a diverse and complex genus (Pisanty et al., 2022). Similarly for the enormous genus *Lasioglossum* (Halictidae: Halictinae). The position of the subfamily Anthophorinae (Apidae) is also uncertain, having been recovered as sister to Nomadinae (Bossert et al., 2019), as well as sister to all remaining Apidae families (Orr et al., 2022). Our all-species tree agrees with the latter but with weak support (ufbs 73%) while our genus tree has yet another result (Fig.S1). A similar case applies with the mono- or paraphyly of the Centridini. We summarise key differences between this phylogeny and previously published work in Supplementary file 5, which also mentions some genera that are potentially not monophyletic.

### 3.5. Dating: keeping it simple

For large phylogenies, Bayesian dating methods are computationally intensive. Penalised likelihood rate smoothing (PLRS) as implemented in TreePL, is an efficient alternative without the complexities of the divide and graft approach (e.g., Jetz et al., 2012; Smirnov and Warnow, 2021; Upham et al., 2019). We took the simplest dating approach by allowing the data and relaxed-clock model to define the relative age pattern, applying a single overall root calibration to scale the absolute age, the single most important calibration in relaxed-clock dating (Kawahara et al., 2023; Mello and Schrago, 2014; Sanderson, 2002). This result was then used to compare how well the broad tree shape matches previous dating efforts and proposed fossil-based internal calibrations (Beaulieu and O’Meara, 2018; Hugall et al., 2007; Near et al., 2005).

We compared a range of groups across families, for which there were multiple previous estimates from 19 different published studies (Fig. S3A; Supplementary Table S7). We also looked at the 34 fossil calibrations in Cardinal et al. (2018), including 15 previously described in detail in Cardinal and Danforth (2013) (Fig. S3B). Our dating falls within the range for most taxa (Fig. S3A), representing a broad consensus framework, and the all-species and genus only tree dates are highly correlated (0.994). Most of our dates fall either within the Cardinal et al. (2018) prior 95% confidence intervals (CI) or above their minimum age constraint (Fig. S3B). These comparisons highlight the limited knowledge of the age of bees as a whole or the age of many of the lineages. In addition, many studies use previous results as secondary calibrations, meaning there is even less known than it appears. General consilience of tree and fossil constraints suggest that the data+model is reasonable for estimating a time tree. The unconstrained tree is sufficiently clock-like that apparent rate variation can be accommodated by the PLRS correlated model (e.g., Fig. S1).

The fossil constraints are either much younger than the lineage appears to be, or they are close. As they are essentially minimum age constraints (and would only be applied as such in PLRS), they would have limited influence (Kawahara et al., 2023). There are a few exceptions worth noting where we found significant disparities to the Cardinal et al. (2018) fossil calibrations (Table 3, Fig. S3B).

**Table 3.** Differences with Cardinal et al. (2018) fossil calibrations.

|  | Notes |
| --- | --- |
| Andrenidae 14, 19, 20 | Dates for Andreninae and Pangurninae are older, but our taxon sampling is much greater and our Andrenidae dates fall within the range of previous studies (Fig. S3A). Cardinal et al. (2018) dates are at or outside their prior maximum CI. |
| Apidae 3, 24 | Given that the genus *Apis* is not sister to the Euglossini tribe (Bossert et al., 2019), the stem for Apidae might be deeper. In Cardinal et al. (2018) the dating for Apidae is at the maximum of their prior CI. In the case of Melectini, our dating is older but in Cardinal et al. (2018) posterior is at their prior CI maximum. |
| Apidae 4, 5 | These involve the stingless bees (Meliponini). Our dates are younger, and the larger taxon sampling reveals the complexity of genera involved. The whole group appears to have a shallow crown and a long stem. This could be the result of using fossils that are stem taxa and not extant crown taxa, an uncertainty posited for many of the Cardinal et al. (2018) calibrations. |
| Melittidae 33 | Our date for this family is older but considering it is sister to all other families there is scope for a deeper age. |
| Root (crown age) of the extant Anthophila 17 | Cardinal et al. (2018) based this on the fossil *Melittosphex burmensis* but a recent paper suggests it might not be a bee (Rosa and Melo, 2021). Previous estimates, including large hymenopteran phylogenomic studies, returned a wide range of ages. Thus, we kept to a broad consensus normal distribution age (mean 100 mya, SD 4) for the extant Anthophila, minimizing the disparity. |

### 3.6. Direction of future efforts

Considering the amount of UCE data that has become available in recent years, we anticipate large projects that will consolidate all primary UCE. With these data it would be possible to build a more comprehensive bee backbone ‘stub’. In the case of data structure commonality, a targeted approach would be the best way to overcome the current gaps such as the recent targeted exon capture method used to build a global phylogeny for butterflies (Kawahara et al., 2023). Gaps in bee molecular data that emerged from this work are outlined in our summaries of data patchiness. At present most genera are represented but within genera, species sampling is uneven, ranging between zero to 100% but often below 20%. A few key ‘legacy genes’ would be needed, at least initially, such as COI and EF-1α. Substantial mtDNA often comes with raw UCE data (‘genome skimming’; e.g., Branstetter et al. (2021), Sann et al. (2021)), so it is worthwhile including this in order to link species with UCE to the large COI barcode database.

It is important to recognise the limitations of this phylogeny. For most bee species, there is no molecular data available and thus, their placement and relationships remain uncertain. Studies focusing on patterns of diversification in space and time have overcome incomplete sampling by using the birth–death polytomy resolvers of Kuhn et al. (2011) and Thomas et al. (2013), which assign a position to species with no molecular data into phylogenetic trees using information based on taxonomy. This approach has been used to simulate the phylogenetic positions of taxa in mammals and birds (Jetz et al., 2012; Rabosky et al., 2018; Upham et al., 2019), and while the resulting phylogenies might be reliable for diversification and phylogenetic distinctiveness analyses (but see Chang et al., 2020), they should be used with caution in phylogenetic comparative analyses because the placement of these taxa within a phylogeny does not take into account their species trait values (Rabosky, 2015). Our supermatrix only includes species with molecular data, accounting for 23% of the total number of bee species currently accepted. This is far from the coverage of molecular data available for birds or marine fishes (Jetz et al., 2012; Rabosky et al., 2018), but close to the proportion of seed plants with molecular data (Smith and Brown, 2018). For comparative analyses this supermatrix phylogeny provides enough species for multiple phylogenetically independent comparisons (Supplementary Table S4). Nonetheless, we stress the need for the use of approaches that deal with missing data (Garamszegi and Møller, 2011), and the importance of always assessing the assumptions and biases of phylogenetic comparative analyses, as to avoid poor model fits and misinterpretation of results (Cooper et al., 2016; Rangel et al., 2015). Future efforts should be placed in sampling species from realms, mainly from the Global South, that currently have a limited amount of molecular data available.

## 4. Conclusions

This work presents the most extensive bee phylogeny to date, providing a summary of substantial existing sequence data available up to April 2023, as well as its gaps. The supermatrix phylogeny was built with public data curated with some care, yielding a composite of previously published phylogenetic hypotheses regarding bees that can be downloaded and subsetted online at beetreeoflife.org. Phylogenetic trees in this work show sensible support between bee families, most subfamilies and tribes. At genus level, genera are for the most part represented, but within genera low data coverage remains a problem in some taxa. Future additional work could improve resolution, particularly for genera that remain under-sampled. This in turn could provide a better understanding of the evolution of this diverse group of insects.

### CRediT authorship contribution statement

**Patricia Henríquez-Piskulich:** Conceptualization, Formal analysis, Investigation, Data Curation, Writing - original draft, Writing - review & editing, Visualization. **Andrew F. Hugall:** Conceptualization, Methodology, Formal analysis, Investigation, Data Curation, Writing - original draft, Writing - review & editing, Visualization. **Devi Stuart-Fox:** Conceptualization, Writing - review & editing, Supervision, Funding acquisition.

## Acknowledgments

Patricia Henríquez-Piskulich was supported by funding awarded from the Agencia Nacional de Investigación y Desarrollo de Chile (Scholarship ID 72210037).

## Data availability

All data and supplementary files generated for this study have been deposited at the Dryad Digital Repository (https://doi.org/10.5061/dryad.80gb5mkw1).

