## Supplementary data for "A supermatrix phylogeny of the world’s bees (Hymenoptera: Anthophila)"

**Table S1.** Molecular phylogenetic studies previously published for bees.

| **Study** |
|---|
| Abrahamczyk, S., de Vos, J.M., Sedivy, C., Gottleuber, P., Kessler, M., 2014. A humped latitudinal phylogenetic diversity pattern of orchid bees (Hymenoptera: Apidae: Euglossini) in western Amazonia: Assessing the influence of climate and geologic history. Ecography (Cop.). 37, 500–508. https://doi.org/10.1111/j.1600-0587.2013.00417.x |
| Aguiar, A.J.C., Melo, G.A.R., Vasconcelos, T.N.C., Gonçalves, R.B., Giugliano, L., Martins, A.C., 2020. Biogeography and early diversification of Tapinotaspidini oil-bees support presence of Paleocene savannas in South America. Mol. Phylogenet. Evol. 143, 106692. https://doi.org/10.1016/j.ympev.2019.106692 |
| Almeida, E.A.B., Danforth, B.N., 2009. Phylogeny of colletid bees (Hymenoptera: Colletidae) inferred from four nuclear genes. Mol. Phylogenet. Evol. 50, 290–309. https://doi.org/10.1016/j.ympev.2008.09.028 |
| Almeida, E.A.B., Packer, L., Melo, G.A.R., Danforth, B.N., Cardinal, S.C., Quinteiro, F.B., Pie, M.R., 2019. The diversification of neopasiphaeine bees during the Cenozoic (Hymenoptera: Colletidae). Zool. Scr. 48, 226–242. https://doi.org/10.1111/zsc.12333 |
| Almeida, E.A.B., Pie, M.R., Brady, S.G., Danforth, B.N., 2012. Biogeography and diversification of colletid bees (Hymenoptera: Colletidae): Emerging patterns from the southern end of the world. J. Biogeogr. 39, 526–544. https://doi.org/10.1111/j.1365-2699.2011.02624.x |
| Arias, M.C., Sheppard, W.S., 2005. Phylogenetic relationships of honey bees (Hymenoptera:Apinae:Apini) inferred from nuclear and mitochondrial DNA sequence data. Mol. Phylogenet. Evol. 37, 25–35. https://doi.org/10.1016/j.ympev.2005.02.017 |
| Ascher, J.S., 2004. Systematics of the bee family Andrenidae (Hymenoptera: Apoidea). ProQuest Dissertations Publishing. |
| Ascher, J.S., Danforth, B.N., Ji, S., 2001. Phylogenetic utility of the major opsin in bees (Hymenoptera: Apoidea): A reassessment. Mol. Phylogenet. Evol. 19, 76–93. https://doi.org/10.1006/mpev.2001.0911 |
| Bossert, S., Murray, E.A., Almeida, E.A.B., Brady, S.G., Blaimer, B.B., Danforth, B.N., 2019. Combining transcriptomes and ultraconserved elements to illuminate the phylogeny of Apidae. Mol. Phylogenet. Evol. 130, 121–131. https://doi.org/10.1016/j.ympev.2018.10.012 |
| Bossert, S., Murray, E.A., Pauly, A., Chernyshov, K., Brady, S.G., Danforth, B.N., 2021. Gene Tree Estimation Error with Ultraconserved Elements: An Empirical Study on Pseudapis Bees . Syst. Biol. 70, 803–821. https://doi.org/10.1093/sysbio/syaa097 |
| Bossert, S., Wood, T.J., Patiny, S., Michez, D., Almeida, E.A.B., Minckley, R.L., Packer, L., Neff, J.L., Copeland, R.S., Straka, J., Pauly, A., Griswold, T., Brady, S.G., Danforth, B.N., Murray, E.A., 2022. Phylogeny, biogeography and diversification of the mining bee family Andrenidae. Syst. Entomol. 47, 283–302. https://doi.org/10.1111/syen.12530 |
| Brady, S.G., Litman, J.R., Danforth, B.N., 2011. Rooting phylogenies using gene duplications: An empirical example from the bees (Apoidea). Mol. Phylogenet. Evol. 60, 295–304. https://doi.org/10.1016/j.ympev.2011.05.002 |
| Bull, N.J., Schwarz, M.P., Cooper, S.J.B., 2003. Phylogenetic divergence of the Australian allodapine bees (Hymenoptera: Apidae). Mol. Phylogenet. Evol. 27, 212–222. https://doi.org/10.1016/S1055-7903(02)00402-5 |
| Cameron, S.A., Hines, H.M., Williams, P.H., 2007. A comprehensive phylogeny of the bumble bees (Bombus). Biol. J. Linn. Soc. 91, 161–188. https://doi.org/10.1111/j.1095-8312.2007.00784.x |
| Cameron, S.A., Mardulyn, P., 2001. Multiple molecular data sets suggest independent origins of highly eusocial behavior in bees (Hymenoptera:Apinae). Syst. Biol. 50, 194–214. https://doi.org/10.1080/10635150120230 |

Cameron, S.A., Williams, P.H., 2003. Phylogeny of bumble bees in the New World subgenus Fervidobombus (Hymenoptera: Apidae): Congruence of molecular and morphological data. Mol. Phylogenet. Evol. 28, 552–563. https://doi.org/10.1016/S1055-7903(03)00056-3

Cardinal, S., Danforth, B.N., 2013. Bees diversified in the age of eudicots. Proc. R. Soc. B Biol. Sci. 280. https://doi.org/10.1098/rspb.2012.2686

Cardinal, S., Straka, J., Danforth, B.N., 2010. Comprehensive phylogeny of apid bees reveals the evolutionary origins and antiquity of cleptoparasitism. Proc. Natl. Acad. Sci. U. S. A. 107, 16207–16211. https://doi.org/10.1073/pnas

Chenoweth, L.B., Schwarz, M.P., 2011. Biogeographical origins and diversification of the exoneurine allodapine bees of Australia (Hymenoptera, Apidae). J. Biogeogr. 38, 1471–1483. https://doi.org/10.1111/j.1365-2699.2011.02488.x

Costa, M.A., Del Lama, M.A., Melo, G.A.R., Sheppard, W.S., 2003. Molecular phylogeny of the stingless bees (Apidae, Apinae, Meliponini) inferred from mitochondrial 16S rDNA sequences. Apidologie 34, 73–84.

Danforth, B.N., 1999. Phylogeny of the bee genus Lasioglossum (Hymenoptera: Halictidae) based on mitochondrial COI sequence data. Syst. Entomol. 24, 377–393. https://doi.org/10.1046/j.1365-3113.1999.00087.x

Danforth, B.N., 2002. Evolution of sociality in a primitively eusocial lineage of bees. Proc. Natl. Acad. Sci. U. S. A. 99, 286–290. https://doi.org/10.1073/pnas.012387999

Danforth, B.N., Brady, S.G., Sipes, S.D., Pearson, A., 2004. Single-copy nuclear genes recover cretaceous-age divergences in bees. Syst. Biol. 53, 309–326. https://doi.org/10.1080/10635150490423737

Danforth, B.N., Conway, L., Ji, S., 2003. Phylogeny of eusocial Lasioglossum reveals multiple losses of eusociality within a primitively eusocial clade of bees (Hymenoptera: Halictidae). Syst. Biol. 52, 23–36. https://doi.org/10.1080/10635150390132687

Danforth, B.N., Eardley, C., Packer, L., Walker, K., Pauly, A., Randrianambinintsoa, F.J., 2008. Phylogeny of Halictidae with an emphasis on endemic African Halictinae. Apidologie 39, 86–101. https://doi.org/10.1051/apido:2008002

Danforth, B.N., Fang, J., Sipes, S., 2006. Analysis of family-level relationships in bees (Hymenoptera: Apiformes) using 28S and two previously unexplored nuclear genes: CAD and RNA polymerase II. Mol. Phylogenet. Evol. 39, 358–372.

Danforth, B.N., Ji, S., 2001. Australian Lasioglossum + Homalictus form a monophyletic group: Resolving the "Australian Enigma." Syst. Biol. 50, 268–283. https://doi.org/10.1093/sysbio/50.2.268

Danforth, B.N., Sauquet, H., Packer, L., 1999. Phylogeny of the Bee Genus Halictus (Hymenoptera: Halictidae) Based on Parsimony and Likelihood Analyses of Nuclear EF-1α Sequence Data. Mol. Phylogenet. Evol. 13, 605–618. https://doi.org/10.1006/mpev.1999.0670

Danforth, B.N., Sipes, S., Fang, J., Brady, S.G., 2006. The history of early bee diversification based on five genes plus morphology. Proc. Natl. Acad. Sci. U. S. A. 103, 15118–15123. https://doi.org/10.1073/pnas.0604033103

Dellicour, S., Lecocq, T., Kuhlmann, M., Mardulyn, P., Michez, D., 2014. Molecular phylogeny, biogeography, and host plant shifts in the bee genus Melitta (Hymenoptera: Anthophila). Mol. Phylogenet. Evol. 70, 412–419. https://doi.org/10.1016/j.ympev.2013.08.013

Dorchin, A., Danforth, B.N., Griswold, T., Sinclair, B., Klass, K.-D., 2018. A new genus of eucerine bees endemic to southwestern North America revealed in phylogenetic analyses of the Eucera complex (Hymenoptera: Apidae: Eucerini). Arthropod Syst. Phylogeny 76, 215–234.

Dorchin, A., López-Uribe, M.M., Praz, C.J., Griswold, T., Danforth, B.N., 2018. Phylogeny, new generic-level classification, and historical biogeography of the Eucera complex (Hymenoptera: Apidae). Mol. Phylogenet. Evol. 119, 81–92. https://doi.org/10.1016/j.ympev.2017.10.007

Duennes, M.A., Lozier, J.D., Hines, H.M., Cameron, S.A., 2012. Geographical patterns of genetic divergence in the widespread Mesoamerican bumble bee Bombus ephippiatus (Hymenoptera: Apidae). Mol. Phylogenet. Evol. 64, 219–231. https://doi.org/10.1016/j.ympev.2012.03.018

Ferrari, R.R., Onuferko, T.M., Monckton, S.K., Packer, L., 2020. The evolutionary history of the cellophane bee genus Colletes Latreille (Hymenoptera: Colletidae): Molecular phylogeny, biogeography and implications for a global infrageneric classification. Mol. Phylogenet. Evol. 146, 106750. https://doi.org/10.1016/j.ympev.2020.106750

Freitas, F. V., Branstetter, M.G., Griswold, T., Almeida, E.A.B., 2020. Partitioned Gene-Tree Analyses and Gene-Based Topology Testing Help Resolve Incongruence in a Phylogenomic Study of Host-Specialist Bees (Apidae: Eucerinae). Mol. Biol. Evol. 38, 1090–1100. https://doi.org/10.1093/molbev/msaa277

Freitas, F. V., Santos Júnior, J.E. Dos, Santos, F.R., Silveira, F.A., 2019. A phylogenetic study of the Thygater-Trichocerapis group and new scopes for the subgenera of Thygater Holmberg, (Hymenoptera, Apidae). Syst. Entomol. 44, 728–744. https://doi.org/10.1111/syen.12351

Freitas, F.V., Santos Júnior, J.E., Santos, F.R., Silveira, F.A., 2018. Species delimitation and sex associations in the bee genus Thygater, with the aid of molecular data, and the description of a new species. Apidologie 49, 484–496. https://doi.org/10.1007/s13592-018-0576-0

Gerth, M., Röthe, J., Bleidorn, C., 2013. Tracing horizontal Wolbachia movements among bees (Anthophila): A combined approach using multilocus sequence typing data and host phylogeny. Mol. Ecol. 22, 6149–6162. https://doi.org/10.1111/mec.12549

Gibbs, J., Brady, S.G., Kanda, K., Danforth, B.N., 2012. Phylogeny of halictine bees supports a shared origin of eusociality for Halictus and Lasioglossum (Apoidea: Anthophila: Halictidae). Mol. Phylogenet. Evol. 65, 926–939. https://doi.org/10.1016/j.ympev.2012.08.013

Gonçalves, R.B., 2016. A molecular and morphological phylogeny of the extant Augochlorini (Hymenoptera, Apoidea) with comments on implications for biogeography. Syst. Entomol. 41, 430–440. https://doi.org/10.1111/syen.12166

González-Vaquero, R.A., Roig-Alsina, A., Packer, L., 2016. DNA barcoding as a useful tool in the systematic study of wild bees of the tribe Augochlorini (Hymenoptera: Halictidae). Genome 59, 889–898. https://doi.org/10.1139/gen-2016-0006

Habermannová, J., Bogusch, P., Straka, J., 2013. Flexible Host Choice and Common Host Switches in the Evolution of Generalist and Specialist Cuckoo Bees (Anthophila: Sphecodes). PLoS One 8. https://doi.org/10.1371/journal.pone.0064537

Haider, M., Dorn, S., Sedivy, C., Müller, A., 2014. Phylogeny and floral hosts of a predominantly pollen generalist group of mason bees (Megachilidae: Osmiini). Biol. J. Linn. Soc. 111, 78–91. https://doi.org/10.1111/bij.12186

He, B., Su, T., Niu, Z., Zhou, Z., Gu, Z., Huang, D., 2019. Characterization of mitochondrial genomes of three Andrena bees (Apoidea: Andrenidae) and insights into the phylogenetics. Int. J. Biol. Macromol. 127, 118–125. https://doi.org/10.1016/j.ijbiomac.2019.01.036

Hedtke, S.M., Patiny, S., Danforth, B.N., 2013. The bee tree of life: A supermatrix approach to apoid phylogeny and biogeography. BMC Evol. Biol. 13. https://doi.org/10.1186/1471-2148-13-138

Hines, H.M., Cameron, S.A., 2010. The phylogenetic position of the bumble bee inquiline Bombus inexspectatus and implications for the evolution of social parasitism. Insectes Soc. 57, 379–383. https://doi.org/10.1007/s00040-010-0094-1

Hines, H.M., Cameron, S.A., Williams, P.H., 2006. Molecular phylogeny of the bumble bee subgenus. Invertebr. Syst. 20, 289–303.

Husemann, M., Neiber, M.T., Nickel, J., Reinbold, C.V.M., Kuhlmann, M., Cordellier, M., 2021. Mitogenomic phylogeny of bee families confirms the basal position and monophyly of Melittidae. Zool. Scr. 50, 352–357. https://doi.org/10.1111/zsc.12468

Kahnt, B., Montgomery, G.A., Murray, E., Kuhlmann, M., Pauw, A., Michez, D., Paxton, R.J., Danforth, B.N., 2017. Playing with extremes: Origins and evolution of exaggerated female forelegs in South African Rediviva bees. Mol. Phylogenet. Evol. 115, 95–105. https://doi.org/10.1016/j.ympev.2017.07.025

Kawakita, A., Ascher, J.S., Sota, T., Kato, M., Roubik, D.W., 2008. Phylogenetic analysis of the corbiculate bee tribes based on 12 nuclear protein-coding genes (Hymenoptera: Apoidea: Apidae). Apidologie 39, 163–175. https://doi.org/10.1051/apido:2007046

Kawakita, A., Sota, T., Ascher, J.S., Ito, M., Tanaka, H., Kato, M., 2003. Evolution and phylogenetic utility of alignment gaps within intron sequences of three nuclear genes in bumble bees (Bombus). Mol. Biol. Evol. 20, 87–92. https://doi.org/10.1093/molbev/msg007

Kayaalp, P., Schwarz, M.P., Stevens, M.I., 2013. Rapid diversification in Australia and two dispersals out of Australia in the globally distributed bee genus, Hylaeus (Colletidae: Hylaeinae). Mol. Phylogenet. Evol. 66, 668–678. https://doi.org/10.1016/j.ympev.2012.10.018

Kuhlmann, M., Almeida, E.A.B., Laurenne, N., Quicke, D.L.J., 2009. Molecular phylogeny and historical biogeography of the bee genus Colletes Latreille, 1802 (Hymenoptera: Apiformes: Colletidae), based on mitochondrial COI and nuclear 28s sequence data. Insect Syst. Evol. 40, 291–318. https://doi.org/10.1163/139956009X12475840653733


Kuhlmann, M., Almeida, E.A.B., Laurenne, N., Quicke, D.L.J., 2009. Molecular phylogeny and historical biogeography of the bee genus Colletes Latreille, 1802 (Hymenoptera: Apiformes: Colletidae), based on mitochondrial COI and nuclear 28s sequence data. Insect Syst. Evol. 40, 291–318. https://doi.org/10.1163/139956009X12475840653733


Larkin, L.L., Neff, J.L., Simpson, B.B., 2006. Phylogeny of the Callandrena subgenus of Andrena (Hymenoptera: Andrenidae) based on mitochondrial and nuclear DNA data: Polyphyly and convergent evolution. Mol. Phylogenet. Evol. 38, 330–343. https://doi.org/10.1016/j.ympev.2005.10.003

Leijs, R., Batley, M., Hogendoorn, K., 2017. The genus Amegilla (Hymenoptera, Apidae, Anthophorini) in Australia: A revision of the subgenera Notomegilla and Zonamegilla. Zookeys 653, 79–140. https://doi.org/10.3897/zookeys.653.11177

Leijs, R., Dorey, J., Hogendoorn, K., 2020. The genus Amegilla (Hymenoptera, Apidae, Anthophorini) in Australia: A revision of the subgenus Asaropoda. Zookeys 2020, 45–122. https://doi.org/10.3897/zookeys.908.47375

Leys, R., Cooper, S.J.B., Schwarz, M.P., 2000. Molecular phylogeny of the large carpenter bees, genus Xylocopa (Hymenoptera: Apidae), based on mitochondrial DNA sequences. Mol. Phylogenet. Evol. 17, 407–418. https://doi.org/10.1006/mpev.2000.0851

Leys, R., Hogendoorn, K., 2008. Correlated evolution of mating behaviour and morphology in large carpenter bees (Xylocopa). Apidologie 39, 119–132.

Lim, K., Lee, Seunghyun, Orr, M., Lee, Seunghwan, 2022. Harrison's rule corroborated for the body size of cleptoparasitic cuckoo bees (Hymenoptera: Apidae: Nomadinae) and their hosts. Sci. Rep. 12, 1–12. https://doi.org/10.1038/s41598-022-14938-9

Litman, J.R., Danforth, B.N., Eardley, C.D., Praz, C.J., 2011. Why do leafcutter bees cut leaves? new insights into the early evolution of bees. Proc. R. Soc. B Biol. Sci. 278, 3593–3600. https://doi.org/10.1098/rspb.2011.0365

Litman, J.R., Griswold, T., Danforth, B.N., 2016. Phylogenetic systematics and a revised generic classification of anthidiine bees (Hymenoptera: Megachilidae). Mol. Phylogenet. Evol. 100, 183–198. https://doi.org/10.1016/j.ympev.2016.03.018

Litman, J.R., Praz, C.J., Danforth, B.N., Griswold, T.L., Cardinal, S., 2013. Origins, Evolution, And Diversification Of Cleptoparasitic Lineages In Long-Tongued Bees. Evolution (N. Y). 67, 2982–2998. https://doi.org/10.1111/evo.12161

Liu, X.W., Chesters, D., Dai, Q.Y., Niu, Z.Q., Beckschäfer, P., Martin, K., Zhu, C.D., 2017. Integrative Profiling of Bee Communities from Habitats of Tropical Southern Yunnan (China). Sci. Rep. 7, 1–14. https://doi.org/10.1038/s41598-017-05262-8

López-Uribe, M.M., Zamudio, K.R., Cardoso, C.F., Danforth, B.N., 2014. Climate, physiological tolerance and sex-biased dispersal shape genetic structure of Neotropical orchid bees. Mol. Ecol. 23, 1874–1890. https://doi.org/10.1111/mec.12689

López-Uribe, M.M., Zamudio, K.R., Cardoso, C.F., Danforth, B.N., 2014. Climate, physiological tolerance and sex-biased dispersal shape genetic structure of Neotropical orchid bees. Mol. Ecol. 23, 1874–1890. https://doi.org/10.1111/mec.12689

Magnacca, K.N., Danforth, B.N., 2007. Low nuclear DNA variation supports a recent origin of Hawaiian Hylaeus bees (Hymenoptera: Colletidae). Mol. Phylogenet. Evol. 43, 908–915. https://doi.org/10.1016/j.ympev.2006.09.004

Mardulyn, P., Cameron, S.A., 1999. The Major Opsin in Bees (Insecta: Hymenoptera): A Promising Nuclear Gene for Higher Level Phylogenetics. Mol. Phylogenet. Evol. 12, 168–176. https://doi.org/10.1006/mpev.1998.0606

Martins, A.C., Luz, D.R., Melo, G.A.R., 2018. Palaeocene origin of the Neotropical lineage of cleptoparasitic bees Ericrocidini-Rhathymini (Hymenoptera, Apidae). Syst. Entomol. 43, 510–521. https://doi.org/10.1111/syen.12286

Martins, A.C., Melo, G.A.R., Renner, S.S., 2014. The corbiculate bees arose from New World oil-collecting bees: Implications for the origin of pollen baskets. Mol. Phylogenet. Evol. 80, 88–94. https://doi.org/10.1016/j.ympev.2014.07.003

Michel-Salzat, A., Cameron, S.A., Oliveira, M.L., 2004. Phylogeny of the orchid bees (Hymenoptera: Apinae: Euglossini): DNA and morphology yield equivalent patterns. Mol. Phylogenet. Evol. 32, 309–323. https://doi.org/10.1016/j.ympev.2003.12.009

Michez, D., Eardley, C., Kuhlmann, M., Timmermann, K., Patiny, Ś., 2010. The bee genera Haplomelitta and Samba (Hymenoptera:Anthophila:Melittidae): Phylogeny, biogeography and host plants. Invertebr. Syst. 24, 327–347. https://doi.org/10.1071/IS10008

Michez, D., Patiny, S., Danforth, B.N., 2009. Phylogeny of the bee family Melittidae (Hymenoptera: Anthophila) based on combined molecular and morphological data. Syst. Entomol. 34, 574–597. https://doi.org/10.1111/j.1365-3113.2009.00479.x

Michez, D., Patiny, S., Danforth, B.N., 2009. Phylogeny of the bee family Melittidae (Hymenoptera: Anthophila) based on combined molecular and morphological data. Syst. Entomol. 34, 574–597. https://doi.org/10.1111/j.1365-3113.2009.00479.x

Murray, E.A., Evanhoe, L., Bossert, S., Geber, M.A., Griswold, T., McCoshum, S.M., 2021. Phylogeny, Phenology, and Foraging Breadth of Ashmeadiella (Hymenoptera: Megachilidae). Insect Syst. Divers. 5. https://doi.org/10.1093/isd/ixab010

Onuferko, T.M., Bogusch, P., Ferrari, R.R., Packer, L., 2019. Phylogeny and biogeography of the cleptoparasitic bee genus Epeolus (Hymenoptera: Apidae) and cophylogenetic analysis with its host bee genus Colletes (Hymenoptera: Colletidae). Mol. Phylogenet. Evol. 141, 106603. https://doi.org/10.1016/j.ympev.2019.106603

Orr, M.C., Branstetter, M.G., Straka, J., Yuan, F., Leijs, R., Zhang, D., Zhou, Q., Zhu, C.-D., 2022. Phylogenomic Interrogation Revives an Overlooked Hypothesis for the Early Evolution of the Bee Family Apidae (Hymenoptera: Apoidea), With a Focus on the Subfamily Anthophorinae. Insect Syst. Divers. 6, 1–15. https://doi.org/10.1093/isd/ixac022

Packer, L., Litman, J., Praz, C.J., 2017. Phylogenetic position of a remarkable new fideliine bee from northern Chile (Hymenoptera: Megachilidae). Syst. Entomol. 42, 473–488. https://doi.org/10.1111/syen.12229

Patiny, S., Michez, D., Danforth, B.N., 2008. Phylogenetic relationships and host-plant evolution within the basal clade of Halictidae (Hymenoptera, Apoidea). Cladistics 24, 255–269. https://doi.org/10.1111/j.1096-0031.2007.00182.x

Peters, R.S., Krogmann, L., Mayer, C., Donath, A., Gunkel, S., Meusemann, K., Kozlov, A., Podsiadlowski, L., Petersen, M., Lanfear, R., Diez, P.A., Heraty, J., Kjer, K.M., Klopfstein, S., Meier, R., Polidori, C., Schmitt, T., Liu, S., Zhou, X., Wappler, T., Rust, J., Misof, B., Niehuis, O., 2017. Evolutionary History of the Hymenoptera. Curr. Biol. 27, 1013–1018. https://doi.org/10.1016/j.cub.2017.01.027

Peters, R.S., Meyer, B., Krogmann, L., Borner, J., Meusemann, K., Schütte, K., Niehuis, O., Misof, B., 2011. The taming of an impossible child: A standardized all-in approach to the phylogeny of

Hymenoptera using public database sequences. BMC Biol. 9, 55. https://doi.org/10.1186/1741-7007-9-55

Pisanty, G., Richter, R., Martin, T., Dettman, J., Cardinal, S., 2022. Molecular phylogeny, historical biogeography and revised classification of andrenine bees (Hymenoptera: Andrenidae). Mol. Phylogenet. Evol. 170, 107151. https://doi.org/10.1016/j.ympev.2021.107151

Policarová, J., Cardinal, S., Martins, A.C., Straka, J., 2019. The role of floral oils in the evolution of apid bees (Hymenoptera: Apidae). Biol. J. Linn. Soc. 128, 486–497. https://doi.org/10.1093/biolinnean/blz099

Porto, D.S., Almeida, E.A.B., 2021. Corbiculate Bees (Hymenoptera: Apidae): Exploring the Limits of Morphological Data to Solve a Hard Phylogenetic Problem. Insect Syst. Divers. 5. https://doi.org/10.1093/isd/ixab008

Praz, C., Müller, A., Genoud, D., 2019. Hidden diversity in European bees: Andrena amieti sp. n., a new Alpine bee species related to Andrena bicolor (Fabricius, 1775) (Hymenoptera, Apoidea, Andrenidae). Alp. Entomol. 3, 11–38. https://doi.org/10.3897/alpento.3.29675

Praz, C.J., Müller, A., Danforth, B.N., Griswold, T.L., Widmer, A., Dorn, S., 2008. Phylogeny and biogeography of bees of the tribe Osmiini (Hymenoptera: Megachilidae). Mol. Phylogenet. Evol. 49, 185–197. https://doi.org/10.1016/j.ympev.2008.07.005

Praz, C.J., Packer, L., 2014. Phylogenetic position of the bee genera Ancyla and Tarsalia (Hymenoptera: Apidae): A remarkable base compositional bias and an early Paleogene geodispersal from North America to the Old World. Mol. Phylogenet. Evol. 81, 258–270. https://doi.org/10.1016/j.ympev.2014.09.003

Ramírez, S.R., Nieh, J.C., Quental, T.B., Roubik, D.W., Imperatriz-Fonseca, V.L., Pierce, N.E., 2010. A molecular phylogeny of the stingless bee genus Melipona (Hymenoptera: Apidae). Mol. Phylogenet. Evol. 56, 519–525. https://doi.org/10.1016/j.ympev.2010.04.026

Ramírez, S.R., Roubik, D.W., Skov, C.E., Pierce, N.E., 2010. Phylogeny, diversification patterns and historical biogeography of euglossine orchid bees (Hymenoptera: Apidae). Biol. J. Linn. Soc. 100, 552–572. https://doi.org/10.1111/j.1095-8312.2010.01440.x

Ramos, K.S., Martins, A.C., Melo, G.A.R., 2022. Evolution of andrenine bees reveals a long and complex history of faunal interchanges through the Americas during the Mesozoic and Cenozoic. Mol. Phylogenet. Evol. 172, 107484. https://doi.org/10.1016/j.ympev.2022.107484

Rasmussen, C., Camargo, J.M.F., 2008. A molecular phylogeny and the evolution of nest architecture and behavior in Trigona s.s. (Hymenoptera: Apidae: Meliponini). Apidologie 39, 102–118. https://doi.org/10.1051/apido:2007051

Rasmussen, C., Cameron, S.A., 2007. A molecular phylogeny of the Old World stingless bees (Hymenoptera: Apidae: Meliponini) and the non-monophyly of the large genus Trigona. Syst. Entomol. 32, 26–39. https://doi.org/10.1111/j.1365-3113.2006.00362.x

Rasmussen, C., Cameron, S.A., 2010. Global stingless bee phylogeny supports ancient divergence, vicariance, and long distance dispersal. Biol. J. Linn. Soc. 99, 206–232. https://doi.org/10.1111/j.1095-8312.2009.01341.x

Rehan, S.M., Chapman, T.W., Craigie, A.I., Richards, M.H., Cooper, S.J.B., Schwarz, M.P., 2010. Molecular phylogeny of the small carpenter bees (Hymenoptera: Apidae: Ceratinini) indicates early and rapid global dispersal. Mol. Phylogenet. Evol. 55, 1042–1054. https://doi.org/10.1016/j.ympev.2010.01.011

Rehan, S.M., Leys, R., Schwarz, M.P., 2012. A mid-cretaceous origin of sociality in xylocopine bees with only two origins of true worker castes indicates severe barriers to eusociality. PLoS One 7. https://doi.org/10.1371/journal.pone.0034690

Rehan, S.M., Leys, R., Schwarz, M.P., 2013. First Evidence for a Massive Extinction Event Affecting Bees Close to the K-T Boundary. PLoS One 8. https://doi.org/10.1371/journal.pone.0076683

Ribeiro, T.M.A., Martins, A.C., Silva, D.P., Aguiar, A.J.C., 2021. Systematics of the oil bee genus Lanthanomelissa (Apidae: Tapinotaspidini) and its implications for the biogeography of South American grasslands. J. Zool. Syst. Evol. Res. 59, 1013–1027. https://doi.org/10.1111/jzs.12472

Rightmyer, M.G., Griswold, T., Brady, S.G., 2013. Phylogeny and systematics of the bee genus Osmia (Hymenoptera: Megachilidae) with emphasis on North American Melanosmia: Subgenera, synonymies and nesting biology revisited. Syst. Entomol. 38, 561–576. https://doi.org/10.1111/syen.12013

Sann, M., Meusemann, K., Niehuis, O., Escalona, H.E., Mokrousov, M., Ohl, M., Pauli, T., Schmid-Egger, C., 2021. Reanalysis of the apoid wasp phylogeny with additional taxa and sequence data confirms the placement of Ammoplanidae as sister to bees. Syst. Entomol. 46, 558–569. https://doi.org/10.1111/syen.12475

Sann, M., Niehuis, O., Peters, R.S., Mayer, C., Kozlov, A., Podsiadlowski, L., Bank, S., Meusemann, K., Misof, B., Bleidorn, C., Ohl, M., 2018. Phylogenomic analysis of Apoidea sheds new light on the sister group of bees. BMC Evol. Biol. 18, 1–15. https://doi.org/10.1186/s12862-018-1155-8

Sayol, F., Collado, M., Garcia-Porta, J., Seid, M.A., Gibbs, J., Agorreta, A., Mauro, D.S., Raemakers, I., Sol, D., Bartomeus, I., 2020. Feeding specialization and longer generation time are associated with relatively larger brains in bees: Brain evolution bees. Proc. R. Soc. B Biol. Sci. 287. https://doi.org/10.1098/rspb.2020.0762

Schaefer, H., Renner, S.S., 2008. A phylogeny of the oil bee tribe Ctenoplectrini (Hymenoptera: Anthophila) based on mitochondrial and nuclear data: Evidence for Early Eocene divergence and repeated out-of-Africa dispersal. Mol. Phylogenet. Evol. 47, 799–811. https://doi.org/10.1016/j.ympev.2008.01.030

Schwarz, M., Fuller, S., Tierney, S., Cooper, S.J., 2006. Molecular phylogenetics of the exoneurine allodapine bees reveal an ancient and puzzling dispersal from Africa to Australia. Syst. Biol. 55, 31–45. https://doi.org/10.1080/10635150500431148

Schwarz, M.P., Tierney, S.M., Cooper, S.J.B., Bull, N.J., 2004. Molecular phylogenetics of the allodapine bee genus Braunsapis: A-T bias and heterogeneous substitution parameters. Mol. Phylogenet. Evol. 32, 110–122. https://doi.org/10.1016/j.ympev.2003.11.017

Sedivy, C., Dorn, S., Müller, A., 2013. Molecular phylogeny of the bee genus Hoplitis (Megachilidae: Osmiini) - how does nesting biology affect biogeography? Zool. J. Linn. Soc. 167, 28–42. https://doi.org/10.1111/j.1096-3642.2012.00876.x

Sedivy, C., Praz, C.J., Müller, A., Widmer, A., Dorn, S., 2008. Patterns of host-plant choice in bees of the genus Chelostoma: The constraint hypothesis of host-range evolution in bees. Evolution (N. Y). 62, 2487–2507. https://doi.org/10.1111/j.1558-5646.2008.00465.x

Sipes, S.D., Wolf, P.G., 2001. Phylogenetic relationships within Diadasia, a group of specialist bees. Mol. Phylogenet. Evol. 19, 144–156. https://doi.org/10.1006/mpev.2001.0914

Sless, T.J.L., Branstetter, M.G., Gillung, J.P., Krichilsky, E.A., Tobin, K.B., Straka, J., Rozen, J.G., Freitas, F. V., Martins, A.C., Bossert, S., Searle, J.B., Danforth, B.N., 2022. Phylogenetic relationships and the evolution of host preferences in the largest clade of brood parasitic bees (Apidae: Nomadinae). Mol. Phylogenet. Evol. 166, 107326. https://doi.org/10.1016/j.ympev.2021.107326

Smith, J.A., Chenoweth, L.B., Tierney, S.M., Schwarz, M.P., 2013. Repeated origins of social parasitism in allodapine bees indicate that the weak form of Emery's rule is widespread, yet sympatric speciation remains highly problematic. Biol. J. Linn. Soc. 109, 320–331. https://doi.org/10.1111/bij.12043

Smith, J.A., Tierney, S.M., Park, Y.C., Fuller, S., Schwarz, M.P., 2007. Origins of social parasitism: The importance of divergence ages in phylogenetic studies. Mol. Phylogenet. Evol. 43, 1131–1137. https://doi.org/10.1016/j.ympev.2006.12.028

Soltani, G.G., Bénon, D., Alvarez, N., Praz, C.J., 2017. When different contact zones tell different stories: Putative ring species in the Megachile concinna species complex (Hymenoptera: Megachilidae). Biol. J. Linn. Soc. 121, 815–832. https://doi.org/10.1093/biolinnean/blx023

Tierney, S.M., Sanjur, O., Grajales, G.G., Santos, L.M., Bermingham, E., Wcislo, W.T., 2012. Photic niche invasions: Phylogenetic history of the dim-light foraging augochlorine bees (Halictidae). Proc. R. Soc. B Biol. Sci. 279, 794–803. https://doi.org/10.1098/rspb.2011.1355

Trunz, V., Packer, L., Vieu, J., Arrigo, N., Praz, C.J., 2016. Comprehensive phylogeny, biogeography and new classification of the diverse bee tribe Megachilini: Can we use DNA barcodes in phylogenies of large genera? Mol. Phylogenet. Evol. 103, 245–259. https://doi.org/10.1016/j.ympev.2016.07.004

Wood, T.J., Patiny, S., Bossert, S., 2022. An unexpected new genus of panurgine bees (Hymenoptera, Andrenidae) from Europe discovered after phylogenomic analysis. J. Hymenopt. Res. 89, 183–210. https://doi.org/10.3897/JHR.89.72083

Wright, K.W., Miller, K.B., Song, H., 2020. A molecular phylogeny of the long-horned bees in the genus Melissodes Latreille (Hymenoptera: Apidae: Eucerinae). Insect Syst. Evol. 52, 428–443. https://doi.org/10.1163/1876312X-bja10015

Zappi, A., 2018. Molecular phylogeny of the stingless bees (Apidae, Apinae, Meliponini) inferred from mitochondrial 16S rDNA sequences. Eur. food Res. Technol. = Zeitschrift fur Leb. und -Forschung. A 244, 118–26. https://doi.org/10.1051/apido

Zhang, D., Niu, Z.Q., Luo, A.R., Orr, M.C., Ferrari, R.R., Jin, J.F., Wu, Q.T., Zhang, F., Zhu, C.D., 2022. Testing the systematic status of Homalictus and Rostrohalictus with weakened cross-vein groups within Halictini (Hymenoptera: Halictidae) using low-coverage whole-genome sequencing. Insect Sci. 29, 1819–1833. https://doi.org/10.1111/1744-7917.13034

**Table S2**. Break-down of genes by family, as proportion of species.

| Gene | Apidae | Andrenidae | Halictidae | Colletidae+* | Megachilidae | Melittidae |
|---|---|---|---|---|---|---|
| ***Nuclear*** | | | | | | |
| Phylogenomic stub | 0.01 | 0.00 | 0.01 | 0.00 | 0.01 | 0.04 |
| UCE stub | 0.16 | 0.56 | 0.04 | 0.00 | 0.00 | 0.00 |
| ArgK | 0.34 | 0.00 | 0.00 | 0.03 | 0.00 | 0.00 |
| CAD | 0.03 | 0.02 | 0.10 | 0.02 | 0.60 | 0.24 |
| EF-1α | 0.54 | 0.18 | 0.39 | 0.34 | 0.52 | 0.65 |
| LW Rh | 0.47 | 0.14 | 0.35 | 0.28 | 0.53 | 0.65 |
| NaK | 0.14 | 0.07 | 0.10 | 0.10 | 0.38 | 0.61 |
| Pol II | 0.33 | 0.07 | 0.11 | 0.08 | 0.05 | 0.63 |
| Wnt-1 | 0.14 | 0.04 | 0.39 | 0.33 | 0.07 | 0.39 |
| 28S rDNA | 0.29 | 0.04 | 0.19 | 0.48 | 0.29 | 0.87 |
| ***Mitochondrial*** | | | | | | |
| 16S rDNA | 0.26 | 0.03 | 0.02 | 0.01 | 0.01 | 0.02 |
| COI | 0.82 | 0.76 | 0.89 | 0.90 | 0.84 | 0.74 |
| Cytb | 0.23 | 0.03 | 0.03 | 0.01 | 0.04 | 0.02 |
| **Species** | 1779 | 644 | 779 | 529 | 838 | 82 |
| **Sequences** | 6708 | 1248 | 2021 | 1364 | 2803 | 399 |

*Colletidae+ includes Stenotritidae.

**Table S3**. Taxon sampling by subfamily.

| Family | Subfamily | Genera | Species | Total species | p-species |
|---|---|---|---|---|---|
| Andrenidae | Andreninae | 6 | 469 | 1567 | 0.30 |
| Andrenidae | Oxaeinae | 4 | 8 | 21 | 0.38 |
| Andrenidae | Panurginae | 33 | 167 | 1276 | 0.13 |
| Apidae | Anthophorinae | 5 | 101 | 730 | 0.14 |
| Apidae | Apinae | 53 | 723 | 1296 | 0.56 |
| Apidae | Eucerinae | 52 | 391 | 1152 | 0.34 |
| Apidae | Nomadinae | 56 | 310 | 1600 | 0.19 |
| Apidae | Xylocopinae | 18 | 254 | 1043 | 0.24 |
| Colletidae | Callomelittinae | 1 | 1 | 11 | 0.09 |
| Colletidae | Colletinae | 4 | 137 | 541 | 0.25 |
| Colletidae | Diphaglossinae | 11 | 22 | 136 | 0.16 |
| Colletidae | Euryglossinae | 8 | 21 | 237 | 0.09 |
| Colletidae | Hylaeinae | 5 | 168 | 963 | 0.17 |
| Colletidae | Neopasiphaeinae | 30 | 99 | 414 | 0.24 |
| Colletidae | Scrapterinae | 1 | 17 | 60 | 0.28 |
| Colletidae | Xeromelissinae | 4 | 58 | 135 | 0.43 |
| Halictidae | Halictinae | 41 | 677 | 3323 | 0.20 |
| Halictidae | Nomiinae | 9 | 57 | 576 | 0.10 |
| Halictidae | Nomioidinae | 3 | 6 | 97 | 0.06 |
| Halictidae | Rophitinae | 9 | 39 | 248 | 0.16 |
| Megachilidae | Fideliinae | 4 | 14 | 21 | 0.67 |
| Megachilidae | Lithurginae | 4 | 11 | 59 | 0.19 |
| Megachilidae | Megachilinae | 52 | 811 | 3992 | 0.20 |
| Megachilidae | Pararhophitinae | 1 | 2 | 3 | 0.67 |
| Melittidae | Dasypodainae | 5 | 30 | 93 | 0.32 |
| Melittidae | Meganomiinae | 1 | 1 | 4 | 0.25 |
| Melittidae | Melittinae | 5 | 51 | 100 | 0.51 |
| Stenotritidae | Stenotritinae* | 2 | 6 | 21 | 0.29 |

*Stenotritinae is a default informal name.

**p-species:** proportion of species included in the phylogeny.

**Table S4**. Taxonomic sampling by ecological realm (as defined by Dinerstein et al. (2017)).

| Realm | Species |
|---|---|
| Afrotropic | 370 |
| Australasia | 369 |
| Indomalayan | 283 |
| Nearctic | 1498 |
| Neotropic | 1367 |
| Oceania | 87 |
| Palearctic | 1482 |

**Table S5.** Summary of data structure: genes in common with and between genera, by family.

| Comparison | gic | Andrenidae | Apidae | Colletidae+ | Halictidae | Megachilidae | Melittidae |
|---|---|---|---|---|---|---|---|
| number of genera | | 43 | 184 | 66 | 62 | 61 | 11 |
| within genera | zero | 0.13 | 0.06 | 0.04 | 0.07 | 0.08 | 0.03 |
| | >1 | 0.15 | 0.52 | 0.43 | 0.41 | 0.58 | 0.73 |
| between genera | zero | 0 | 0.09 | 0.02 | 0.12 | 0.01 | 0 |
| | >1 | 0.62 | 0.69 | 0.85 | 0.76 | 0.93 | 1 |
| | median | 2 | 3 | 5 | 4 | 5 | 7 |
| | trip0 | 0 | 0.22 | 0.05 | 0.23 | 0.03 | 0 |
| | trip>1 | 0.46 | 0.53 | 0.78 | 0.64 | 0.90 | 1 |

**gic**: mean number of genes in common for all pairwise taxon comparisons; **trip** is equivalent for taxon triplets (Sanderson et al. 2010).
**within genera**: mean of the genus values for pairwise species comparisons with each genus.
**between genera**: mean of pairwise genus comparisons.
**Colletidae+** includes Stenotritidae.

**Table S6.** Genera with poor data structure by family.

| Family | Genus | species | gic0 | trip0 |
|---|---|---|---|---|
| Apidae | Anthophorula | 13 | 0.38 | 0.58 |
| | Plebeia | 13 | 0.27 | 0.51 |
| | Thyreus | 11 | 0.25 | 0.48 |
| | Triepeolus | 16 | 0.18 | 0.33 |
| | Partamona | 15 | 0.17 | 0.39 |
| | Svastra | 9 | 0.33 | 0.64 |
| Andrenidae | Andrena | 460 | 0.18 | 0.36 |
| | Perdita | 28 | 0.44 | 0.72 |
| | Protandrena | 15 | 0.33 | 0.63 |
| | Calliopsis | 17 | 0.24 | 0.47 |
| | Panurginus | 11 | 0.29 | 0.48 |
| | Macrotera | 9 | 0.42 | 0.75 |
| Colletidae | Hylaeus | 156 | 0.18 | 0.32 |
| | Euhesma | 8 | 0.21 | 0.38 |
| | Caupolicana | 9 | 0.14 | 0.30 |
| Halictidae | Lipotriches | 14 | 0.62 | 0.89 |
| | Nomia | 16 | 0.43 | 0.69 |
| | Dufourea | 18 | 0.16 | 0.29 |
| | Pseudapis | 10 | 0.13 | 0.28 |
| | Augochlora | 8 | 0.43 | 0.75 |
| Megachilidae | Stelis | 40 | 0.29 | 0.55 |
| | Pseudoanthidium | 18 | 0.29 | 0.55 |
| | Anthidium | 41 | 0.15 | 0.26 |
| | Dianthidium | 10 | 0.31 | 0.53 |
| | Heriades | 12 | 0.14 | 0.25 |

**species**: number of species in datamatrix; **gic0**: proportion of pairwise comparisons with no genes in common.
**trip0**: proportion of triplets with no genes in common (Sanderson et al., 2010); gic0 and trip0 are highly correlated.

**Table S7.** References for the dating comparisons.

| Study |
|---|
| Almeida EAB, Pie MR, Brady SG, Danforth BN. 2012. Biogeography and diversification of colletid bees (Hymenoptera: Colletidae): Emerging patterns from the southern end of the world. J Biogeogr. 39(3):526–544. |
| Almeida AB, Packer L, Melo GAR, Danforth BN, Cardinal SC, Quinteiro FB, Pie MR. 2019. The diversification of neopasiphaeine bees during the Cenozoic (Hymenoptera: Colletidae). Zoologica Scripta. 48:226–242. |
| Blaimer BB, Santos BF, A, Gates MW, Kula RR, Mikó I, Rasplus J-Y, Smith DR, Talamas EJ, Brady SB, Buffington ML. 2023. Key innovations and the diversification of Hymenoptera. Nature Communications 14:1212. |
| Bossert S, Wood TJ, Patiny S, Michez D, Almeida EAB, Minckley RL, Packer L, Neff JL, Copeland RS, Straka J, Pauly A, Griswold T, Brady SG, Danforth BN, Murray EA. 2021. Phylogeny, biogeography and diversification of the mining bee family Andrenidae. Systematic Entomology 2022;1–20. |
| Branstetter MG, Danforth BN, Pitts JP, Faircloth BC, Ward PS, Buffngton ML, Gates MW, Kula RR, Brady SG. 2017. Phylogenomic insights into the evolution of stinging wasps and the origins of ants and bees. Curr. Biol. 27: 1019–1025. |
| Cardinal S, Danforth BN. 2013. Bees diversified in the age of eudicots. Proceedings of the Royal Society B: Biological Sciences, 280(1755), 2686. |
| Cardinal S, Buchmann SL, Russell AL. 2018. The evolution of foral sonication, a pollen foraging behavior used by bees (Anthophila). Evolution. 72: 590–600. |
| Freitas FV, Branstetter MG, Griswold T, Almeida EAB. 2021. Partitioned Gene-Tree Analyses and Gene-Based Topology Testing Help Resolve Incongruence in a Phylogenomic Study of Host-Specialist Bees (Apidae: Eucerinae). Mol. Biol. Evol. 38(3):1090–1100. |
| Martins AC, Melo GAR. 2016. The New World oil-collecting bees Centris and Epicharis (Hymenoptera, Apidae): Molecular phylogeny and biogeographic history. Zoologica Scripta, 45(1), 22–33. |
| Martins AC, Luz DR, GAR Melo. 2018. Palaeocene origin of the Neotropical lineage of cleptoparasitic bees Ericrocidini-Rhathymini (Hymenoptera, Apidae). Systematic Entomology (43, 510–521. |
| Odanaka KA, Branstetter MG, Tobin KB, Rehan SM. 2022. Phylogenomics and historical biogeography of the cleptoparasitic bee genus Nomada (Hymenoptera: Apidae) using ultraconserved elements. Molecular Phylogenetics and Evolution 170: 107453. |
| Pisanty G, Richter R, Martin T, Dettman J, Cardinal S. 2022. Molecular phylogeny, historical biogeography and revised classification of andrenine bees (Hymenoptera: Andrenidae). Molecular Phylogenetics and Evolution 170: 107151 |
| Peters RS, Mayer C, Donath A, Gunkel S, Meusemann K, Alexey Kozlov A, Podsiadlowski L, Petersen M, Lanfear R, Diez PA, John Heraty, Karl M. Kjer KM, Klopfstein S, Meier R, Polidori C, Thomas Sch LK. 2017. Evolutionary History of the Hymenoptera. Curr Biol. 7:1013. |
| Ramos KS, Martins AC, Melo GAR. 2022. Evolution of andrenine bees reveals a long and complex history of faunal interchanges through the Americas during the Mesozoic and Cenozoic. Molecular Phylogenetics and Evolution 172: 107484. |
| Rehan SM, Chapman T, Craigie A, Richards MH, Cooper SJB, Schwarz MP. 2010. Molecular phylogeny of the small carpenter bees (Hymenoptera: Apidae: Ceratinini) indicates early and rapid global dispersal. Mol Phylogenet Evol 55: 1042–1054. |
| Rehan SM, Leys R, Schwarz MP. 2012 A Mid-Cretaceous origin of sociality in xylocopine bees with only two origins of true worker caster indicates severe barriers to eusociality. PLoS ONE 7, e34690. |
| Sann M, Niehuis O, Peters RS, Mayer C, Kozlov A, Podsiadlowski L, Bank S, Meusemann K, Misof B, Bleidorn C, et al. 2018. Phylogenomic analysis of Apoidea sheds new light on the sister group of bees. BMC Evol Biol. 18:1–15. |

| |
|---|
| Schwarz MP, Fuller S, Tierney SM, Cooper SJB. 2006. Molecular phylogenetics of the exoneurine allodapine bees reveal an ancient and puzzling dispersal from Africa to Australia. Syst. Biol. 55, 31–45. |
| Trunz V, Packer L, Vieu J, Arrigo N, Praz CJ. 2016. Comprehensive phylogeny, biogeography and new classification of the diverse bee tribe Megachilini: Can we use DNA barcodes in phylogenies of large genera? Molecular Phylogenetics and Evolution 103: 245–259. |

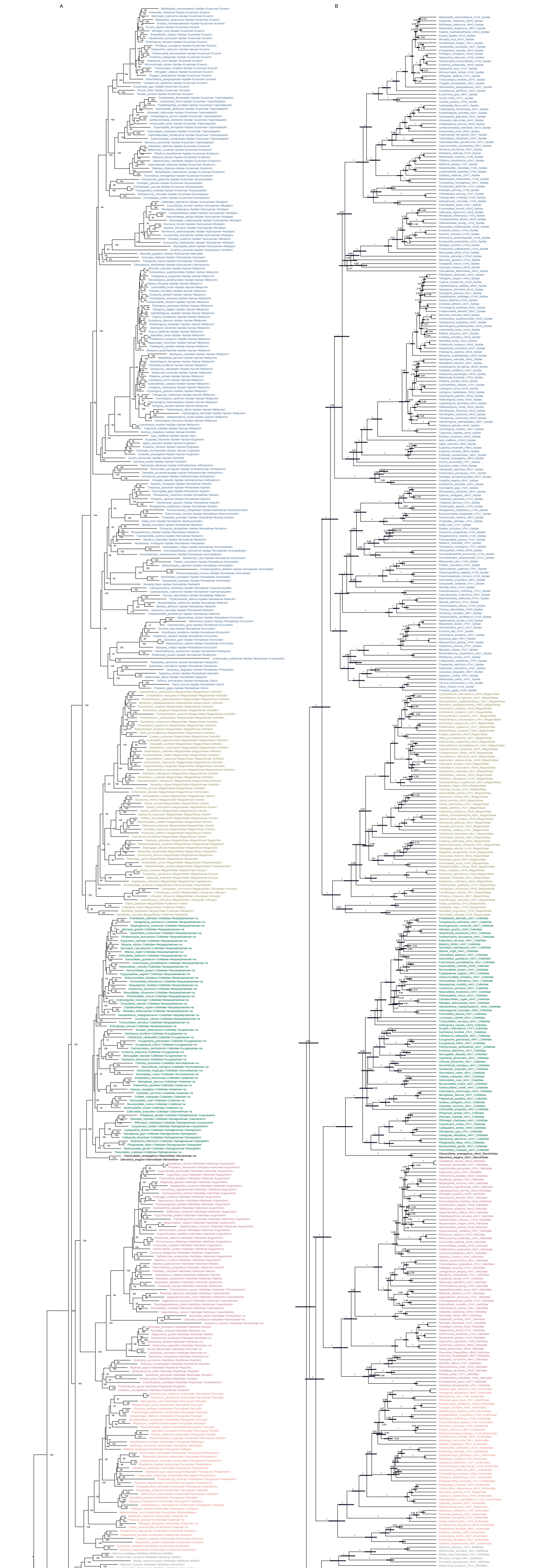


**Figure S1**. Genus representative tree. **A**. IQTree with ufbs percentage node support. Taxon labels indicate genus_species~family~subfami-ly~tribe, coloured by family. Grey 'Crabronidae' indicate outgroups. **B**. Maximum Clade Credibility tree summary of 200 IQTree ufbs trees converted to ultrametric with TreePL penalized likelihood rate smoothing. Labels show genus representative species, number of nuclear and mito-chondrial genes (n#m#) and family. Node support values are indicated as proportion (rather than percentage as in A). Node bars show 95%HPD. Outgroups removed. Summary tree done in BEAST TreeAnnotator.

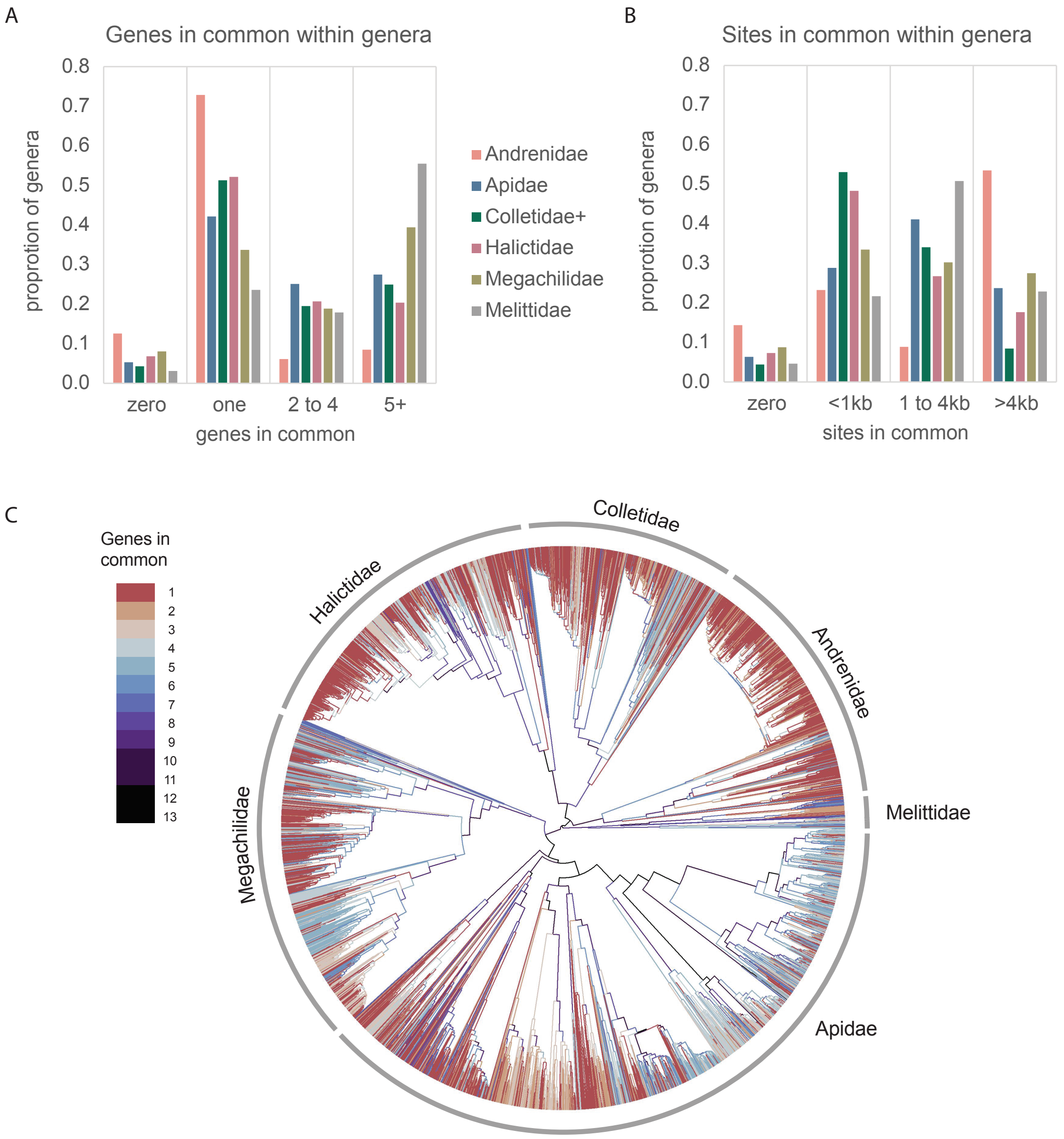


**Figure S2**. Further summaries of data completeness, focussing on amount of overlapping data between groups. **A.** Number of genes in common (gic) between species within genera, by family. The y-axis is the mean across all genera for the proportion of pairs of species that have the given number (x-axis) of genes in common. As in Figure 3, Colletidae+ includes Stenotritidae. **B.** Similar but using sites in common. **C.** Number of genes common to sister clades, with select groups highlighted. Note that for a maximum likelihood tree gic must be >0. See also Tables S2, S5, S6.

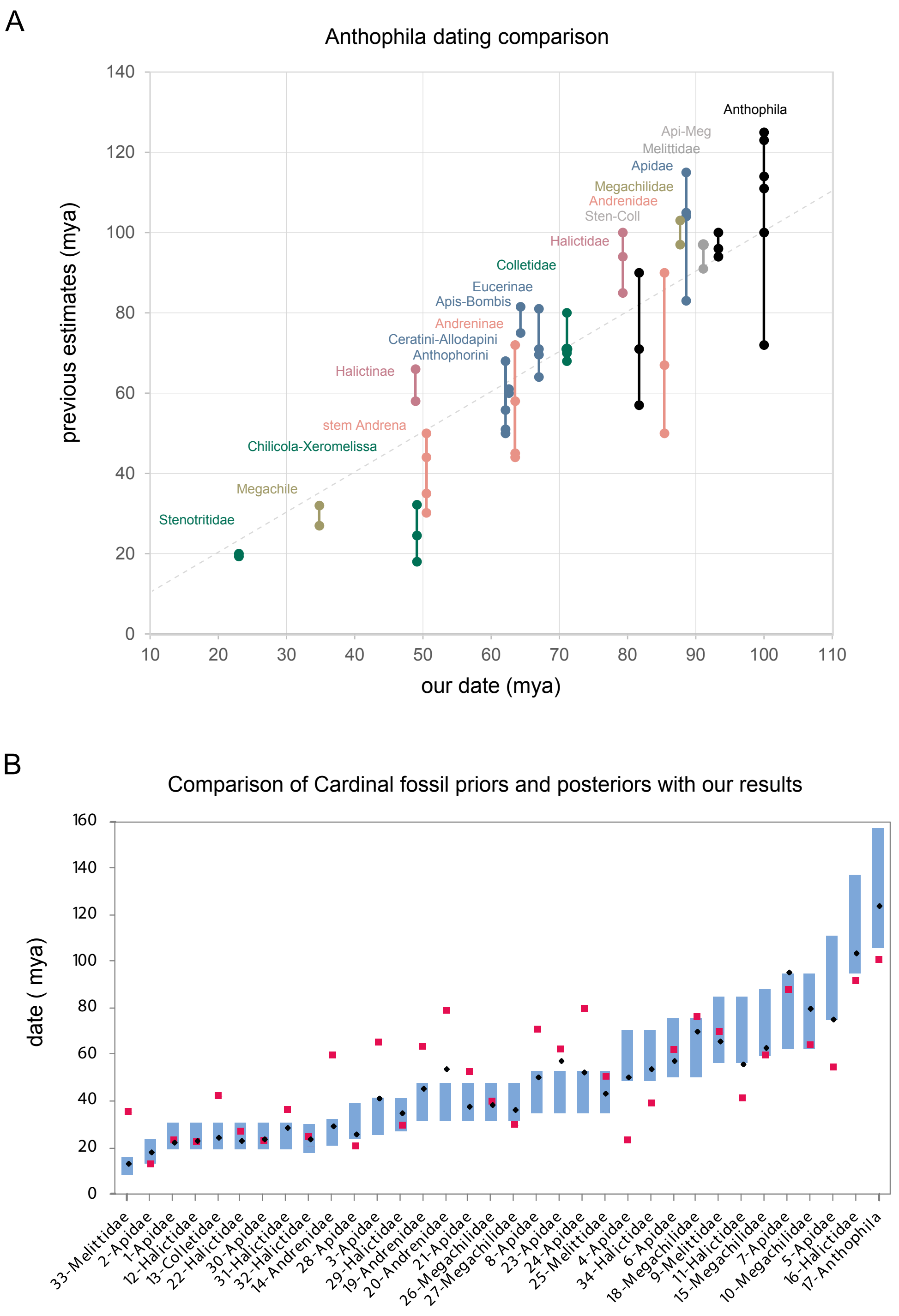


**Figure S3**. Anthophila dating comparisons. **A.** Previous estimates for select groups (y-axis) compared to our results (x-axis). Previous estimates indicated by dots and joined by lines to show the range, coloured by family (see previous figures; super-family groups in grey; dotted line = one-to-one). **B.** Comparison of Cardinal et al. (2018; Table S3) fossil prior 95% HPD range (bars), posterior means (black dots), with our results (red squares). The 34 groups are arranged by Cardinal prior mean and labelled by calibration number and relevant family. See text for details on groups and studies. Using most comparable clades. Sten-Coll = Stenotriti-dae+Colletidae, Api-Meg = Apidae+Megachilidae. Table S9 lists the published studies.
