## Supplementary file 4 for "A supermatrix phylogeny of the world’s bees (Hymenoptera: Anthophila)"

**Distribution of species with molecular data**

To obtain the geographical distribution of bees present in the phylogeny and produce the heat map, we loosely followed the steps described in Orr et al. (2020). We downloaded occurrence data from four public databases (iDigBio (iDB), Global Biodiversity Information Facility (GBIF), Symbiota Collections of Arthropods Network (SCAN), and Atlas of Living Australia (ALA), and collated them using the current most comprehensive curated checklist available (Ascher and Pickering, 2020). All records were downloaded individually and checked for mismatches in the number of records between the website and the databases downloaded. First, we cleaned each database using R 4.2.0 (R Core Team, 2022) and the packages CoordinateCleaner (Zizka et al., 2019), and tidyverse (Wickham et al. 2019). For each database, the automated cleaning steps included removing records that did not have geographical coordinates, coordinates that were not from the country indicated, records with equal absolute longitude and latitude, outlier records for each species, coordinates that fell in the ocean, and coordinates with plain zeros. We then removed records with coordinate precision below 100 Km, records from unsuitable data sources, and duplicated records. In addition, we removed data for species from databases that combined occurrences of two or more species based on synonymy and that were not recognized by the phylogeny to avoid any errors. Then, each database was collated against the Discover Life checklist, and all records that did not match checklist countries (ISO2 code) for each species were removed. The checklist Hierarchical administrative subdivision codes (HASC) were not used to clean the databases as for species with fewer records this approach could be too stringent and remove correct records.

We then compared the final merged database to the taxonomic database (Supplementary file 1) to assess the number of species that remained without distribution data, 389 species did not have records in the public databases downloaded. For these missing species we used the distribution data from the checklist. After this final step, 23 species remained without distribution data.

We produced a heat map for the final clean database merged with the checklist data from a raster file created using R 4.2.0 and the packages sp (Pebesma and Bivand, 2005), sf (Pebesma, 2018), rnaturalearth (Massicotte and South, 2023), rworldmap (South, 2011), and smoothr (Strimas-Mackey, 2018). This raster file and a shapefile of the world map were exported to QGIS to plot the heat maps which were later edited in Adobe Illustrator.

R Core Team, 2022. R: A language and environment for statistical computing.

Symbiota Collections of Arthropods Network (9 March 2023) SCAN Occurrence Download at https://scan-bugs.org/portal/collections/index.php

South, A., 2011. rworldmap: A new R package for mapping global data. R J. 3, 35–43. https://doi.org/10.32614/rj-2011-006

Strimas-Mackey, M., 2018. Smoothr: Smooth and tidy spatial features. R Packag. Version 0.1. 0. Available online https//CRAN. R-project. org/package= smoothr.

Wickham, H., Averick, M., Bryan, J., Chang, W., McGowan, L., François, R., Grolemund, G., Hayes, A., Henry, L., Hester, J., Kuhn, M., Pedersen, T., Miller, E., Bache, S., Müller, K., Ooms, J., Robinson, D., Seidel, D., Spinu, V., Takahashi, K., Vaughan, D., Wilke, C., Woo, K., Yutani, H., 2019. Welcome to the tidyverse. J. Open Source Softw. 4, 1686. https://doi.org/10.21105/joss.01686

Zizka, A., Silvestro, D., Andermann, T., Azevedo, J., Duarte Ritter, C., Edler, D., Farooq, H., Herdean, A., Ariza, M., Scharn, R., Svantesson, S., Wengström, N., Zizka, V., Antonelli, A., 2019. CoordinateCleaner: Standardized cleaning of occurrence records from biological collection databases. Methods Ecol. Evol. 10, 744–751. https://doi.org/10.1111/2041-210X.13152
