## Supplementary file 5 for "A supermatrix phylogeny of the world’s bees (Hymenoptera: Anthophila)"

**Differences between our results and recent work in bee phylogenetics**

**1. Andrenidae**

We recovered most of the relationships previously outlined by Bossert et al. (2021) and Pisanty et al. (2022) with high support, but found a few differences. The genus *Alocandrena* has varied in its position, as either sister to *Andrena* + *Cubiandrena* in Bossert et al. (2021), as well as *Megandrena* + *Ancylandrena* in Pisanty et al. (2022). Our results for both all-species and genus representative trees, showed weak support for the placement of *Alocandrena* as sister to all four, with IQTree ultrafast bootstrap support (ufbs, here and below) 81/67%.

Within Panurginae our results broadly overlap with the findings of Bossert et al. (2021). We found that Perditini, Panurgini, and Protandrenini, were well supported (ufbs 100%). However, relationships within are less so as the position of the genera *Pseudomacrotera*, *Protomeliturga* and *Flavomeliturgula* had low support (59/76/95% respectively) and varied somewhat between all-species and genus trees. *Panurginus* remains paraphyletic as it contains *Clavipanurgus*.

Although we did not focus on relationships at the subgenus level, it is important to highlight that the subgeneric relationships within *Andrena* have also been the focus of recent study. Reclassification changes given its high degree of paraphyly and polyphyly will probably continue to occur as further research is necessary to resolve such diverse and complex genus (Pisanty et al., 2022).

**2. Apidae**

The tribe Anthophorini has recently been raised to its own subfamily, Anthophorinae, and recovered as sister to Nomadinae (Bossert et al., 2019), as well as sister to all remaining Apidae families (Orr et al., 2022). Our result agrees with the former but should be treated with caution as we found weak support with very short internodes (ufbs 73%), and the genus tree returned yet another arrangement (Fig. S1). Within Apinae, the tribe Centridini was monophyletic and remained as the sister group to corbiculates as in Bossert et al. (2019), but in our genus representative tree Centridini was paraphyletic; both trees showed weak support and short internodes (49/73% respectively). Including the data from Freitas et al. (2022) recovered a sister relationship between the genera *Aglae* and *Euglossa* of the tribe Euglossini with short internodes. This differs from the results of Bossert et al. (2019), where the cleptoparasitic genera *Aglae* and *Exaerete* where recovered as sister taxa. Our results mirror this uncertainty, with the genus tree showing *Aglae* as sister to both.

Our results agree with the recent proposal of the subfamily Eucerinae, which includes the tribes Ancylaini, Ancyloscelidini, Emphorini, Eucerini, Exomalopsini, and Tapinotaspidini (Freitas et al., 2020) (ufbs 100% both trees). The tribe Ancyloscelidini has been found to be sister to the Exomalopsini by Freitas et al. (2020), and our results agree but with modest support and short internode (ufbs 83%). The "*Eucera* complex" (Eucerini) has recently undergone a major generic reclassification. *Tetralonia*, *Peponapis*, *Xenoglossa*, *Cemolobu*s, and *Syntrichalonia* have become subgenera of the genus *Eucera,* while *Xenoglossodes* has been re-established as a valid subgenus of *Eucera*, the subgenera *Tetraloniella* synonymized with *Tetralonia*, and *Cubitalia* with *Eucera* (Dorchin et al., 2018b). Furthermore, the venusta-group previously belonging to the *Eucera* subgenus *Synhalonia*, has been recognised as a new genus, *Protohalonia*, a monophyletic group sister to *Simanthedon* (Dorchin et al., 2018a). In our phylogeny, *Eucera* included species belonging to the subgenera Tetralonia, Mirnapis, and Eucera.

All cleptoparasitic genera are a part of the subfamily Nomadinae with the exception of *Ctenoplectrina* (Ctenoplectrini), *Aglae* and *Exaerete* (Bossert et al., 2019). As with Sless et al. (2022), *Ammobates* (Ammobatini) remains paraphyletic due to nested members of the genera *Oreopasites*. As found by Sless et al., (2022), our results recovered a paraphyletic *Melecta* (Melectini) with *Brachymelecta* nested within it. The genus *Coelioxoides* (Coelioxoidini) was one of the longest branches within our results as in previous studies (Orr et al., 2022; Sless et al., 2022), and as such, its placement should be interpreted with caution.

**3. Colletidae and Stenotritidae**

At present there are eight recognized subfamilies (Danforth et al., 2019): Callomelittinae, Colletinae, Diphaglossinae, Euryglossinae, Hylaeinae, Neopasiphaeinae, Scrapterinae, and Xeromelissinae. These subfamilies are mostly well supported but there is negligible phylogenomic data available for Colletidae. We differed in our results with Almeida et al. (2019) only in that we recovered the genus *Scrapter* nested within Euryglossinae. Further taxon sampling may be needed to resolve the relationship between the latter and the subfamily Scrapterinae.

Stenotritidae is a small family of 21 species restricted to Australia, and sister lineage to Colletidae (Danforth et al., 2019). Stenotritidae showed high support in both our full tree and the genera tree, but the two genera described for this family are not monophyletic as seen previously (Almeida et al., 2019).

**4. Halictidae**

Our results recovered subfamily relationships with high support as found in previous work (Danforth et al., 2008, 2004; Patiny et al., 2008). Tribes were for the most part resolved with the exception that *Eupetersia* was nested within the tribe Halictini when in the past it has been placed within Sphecodini (Danforth et al., 2008; Gibbs et al., 2012), and that *Agapostemon* was sister to all remaining Caenohalictini (ufbs 100% all trees) but apparently also contains *Rhinetula denticrus* (94% all trees).

**5. Megachilidae**

There are four recognized subfamilies within Megachilidae and the relationships between them have been assessed through morphological (Gonzalez et al., 2012), and molecular data (Litman et al., 2011; Packer et al., 2017). We found high support for subfamilies and tribes when splitting the genera *Ochreriades*, *Afroheriades* and *Pseudoheriades* into informal monotypic tribes. This was done only for the purposes of our phylogeny, as it is out of the scope of this work to propose formal nomenclature changes.

Our molecular phylogeny differed from the work done in Litman et al. (2011) and Packer et al. (2017), in that it recovered Fideliinae as monophyletic and sister to the other three subfamilies with some support (82% all-species tree).

**6. Melittidae**

Our results agree with previous work which has recovered all the subfamilies, and most tribes and genera as monophyletic, with *Redivivoides* (Melittini) nested within *Rediviva* (Dellicour et al., 2014; Michez et al., 2009).

**Some genera worth re-visiting**

Notwithstanding tree inference limitations and nomenclatural differences, multi-gene data in common identifies a few genera that do not appear to be monophyletic ("modest BS" indicates ufbs <90%, "UCE data" indicates consistent with UCE). We do not claim that these are not monophyletic, only that they worth looking into again; some have already been mentioned in the above sections. Several have been noted before in published studies (e.g., references in Table S1). These genera are:

1. Andrenidae
    a. *Panurginus*: includes *Clavipanurgus*

2. Megachilidae
    a. *Anthodioctes*: (modest BS) in clade with other Anthidiini
    b. *Trichothurgus*: (modest BS) includes *Lithurgopsis*
    c. *Plesianthidium*: (modest BS) Anthidiini complex with *Pachyanthidium*

   d. *Lithurgus*: (modest BS) paraphletic within Lithurgini
   e. *Pachyanthidium*: (modest BS)
   f. *Fidelia*: (modest BS) includes *Fideliopsis*
   g. *Stelis*: includes *Euaspis*

3. Melittidae
   a. *Rediviva*: includes *Redivivoides*

4. Apidae
   a. *Ammobates*: paraphyletic with *Oreopasites* (UCE data)
   b. *Chalepogenus*: several lineages within Tapinotaspidini (UCE data)
   c. *Hasinamelissa*: paraphyletic with *Compsomelissa* complex
   d. *Homotrigona*: complex in Meliponini with *Tetrigona*, *Odontotrigona*
   e. *Frieseomelitta*: complex within Meliponini (albeit limited data)
   f. *Gaesischia*: due to *G. sapucacensis* and *G. mimetica* from subgenus *Dasyhalonia* (UCE data)
   g. *Leurotrigona*: (modest BS) paraphyletic with *Trigonista*
   h. *Melecta*: paraphyletic with *Brachymelecta californica* (UCE data)
   i. *Mesonychium*: includes *Epiclopus gayi*
   j. *Tropidopedia*: (modest BS) paraphyletic with *Lophopedia*

5. Colletidae
   a. *Brachyglossula*: divergent within Neopasiphaeinae
   b. *Caupolicana*: paraphyletic within Caupolicanini
   c. *Edwyniana*: (modest BS) complex within Neopasiphaeinae
   d. *Hylaeus*: (modest BS) may include *Meroglossa*, *Amphylaeus*
   e. *Leioproctus*: complex within Neopasiphaeinae including *Goniocolletes* and others
   f. *Perditomorpha*: (modest BS) complex within Neopasiphaeinae (including *Edwyniana*)
   g. *Ctenocolletes*: paraphyletic with *Stenotritus* sp. (only single sample)

6. Halictidae
   a. *Augochloropsis*: divergent *Augochloropsis semiramis* with *Augochlora*
   b. *Lipotriches*: *Lipotriches australica* with *Nomia*
   c. *Megalopta*: *Megalopta atra* associated with *Xenochlora*
   d. *Ruizantheda*: paraphyletic with *Pseudagapostemon*
